# Leukemic cell-secreted interleukin-9 suppresses cytotoxic T cell-mediated killing in chronic lymphocytic leukemia

**DOI:** 10.1101/2023.10.23.563260

**Authors:** Gioia Boncompagni, Vanessa Tatangelo, Ludovica Lopresti, Cristina Ulivieri, Nagaja Capitani, Carmela Tangredi, Francesca Finetti, Giuseppe Marotta, Federica Frezzato, Andrea Visentin, Sara Ciofini, Alessandro Gozzetti, Monica Bocchia, Diego Calzada-Fraile, Noa B. Martin Cofreces, Livio Trentin, Laura Patrussi, Cosima T. Baldari

**Affiliations:** Department of Life Sciences, University of Siena, Siena, Italy; Stem Cell Transplant and Cellular Therapy Unit, University Hospital, Siena, Italy; Department of Medicine, Hematology and Clinical Immunology Branch, Padua University School of Medicine, Padua, Italy; Venetian Institute of Molecular Medicine, Padua, Italy; Department of Medical Science, Surgery and Neuroscience, University of Siena, Siena, Italy; Immunology Unit from Hospital Universitario de la Princesa, Universidad Autónoma de Madrid and Instituto de investigación Sanitaria La Princesa (IIS-IP), Madrid, Spain; Centro Nacional de Investigaciones Cardiovasculares (CNIC), 28029, Madrid, Spain; Centro de Investigación Biomédica en Red Enfermedades Cardiovasculares (CIBERCV), Madrid, Spain

## Abstract

The tumor microenvironment (TME) plays a central role in the pathogenesis of chronic lymphocytic leukemia (CLL), contributing to disease progression and chemoresistance. Leukemic cells shape the TME into a pro-survival and immunosuppressive niche through contact-dependent and contact-independent interactions with the cellular components of the TME. Immune synapse (IS) formation is defective in CLL. Here we asked whether soluble factors released by CLL cells contribute to their protection from cytotoxic T cell (CTL)-mediated killing by interfering with this process. We found that healthy CTLs cultured in media conditioned by leukemic cells from CLL patients or Eμ-TCL1 mice upregulate the exhaustion marker PD-1 and become unable to form functional ISs and kill target cells. These defects were more pronounced when media were conditioned by leukemic cells lacking p66Shc, a proapoptotic adaptor whose deficiency has been implicated in disease aggressiveness both in CLL and in the Eμ-TCL1 mouse model. Multiplex ELISA assays showed that leukemic cells from Eμ-TCL1 mice secrete abnormally elevated amounts of CCL22, CCL24, IL-9 and IL-10, which are further upregulated in the absence of p66Shc. Among these, IL-9 and IL-10 were also overexpressed in leukemic cells from CLL patients, where they inversely correlated with residual p66Shc. Using neutralizing antibodies or the recombinant cytokines we show that IL-9, but not IL-10, mediates both the enhancement in PD-1 expression and the suppression of effector functions in healthy CTLs. Our results demonstrate that IL-9 secreted by leukemic cells negatively modulates the anti-tumor immune abilities of CTLs, highlighting a new suppressive mechanism and a novel potential therapeutical target in CLL.

## Introduction

To escape immune surveillance, neoplastic B cells of hematologic malignancies implement strategies that shape T lymphocytes of the tumor microenvironment (TME) toward an exhausted phenotype, characterized by increased expression of exhaustion markers and inability to produce adequate levels of immune-activating cytokines, which both contribute to T-cell dysfunctions [1]. T-cell suppression strategies are exploited by tumor cells of chronic lymphocytic leukemia (CLL), the most common leukemia in Western countries. CLL is characterized by the accumulation of mature B cells in peripheral blood and lymphoid organs [2]. Despite a bias of T lymphocytes toward the CD8^+^ phenotype in CLL patients [3,4], the killing activity of cytotoxic T lymphocytes (CTLs), the CD8^+^ effector T cells specialized to eliminate tumor cells [5,6], is defective, mainly as the result of high and sustained expression of inhibitory receptors such as Programmed cell Death protein (PD)-1, Cytotoxic T-Lymphocyte Antigen 4 (CTLA-4) and Lymphocyte-activation gene 3 (LAG-3) [7,8].

The complex mechanisms exploited by T cells to counteract tumor development rely on architectural changes occurring early after antigen recognition that lead to the polarization of receptors, adhesion molecules, cytoskeletal components and organelles toward the T cell/APC contact area to form a highly organized signaling and secretory platform known as the immune synapse (IS) [9,10]. IS formation is a key prerequisite for target cell killing by CTLs [11]. CLL cells disable CTLs through interactions mediated by immunosuppressive surface ligands such as the PD-1 ligand (PD-L1) which, by activating the inhibitory cognate receptors on CTLs, hamper IS formation and polarized lytic granule secretion of into the synaptic cleft to allow for selective tumor cell killing [7,12].

Within the TME, CLL cells secrete cytokines and chemokines which sustain and fuel pro-tumorigenic loops [13–15], as exemplified by Interleukin (IL)-10, that suppresses the ability of monocytes/macrophages to produce TNF-α [16], and IL-9, released by CLL cells from patients with aggressive disease, that stimulates stromal cells to secrete homing chemokines, which in turn contribute to attract CLL cells to the pro-survival lymphoid niche [15,17]. Moreover, IL-9 blockade in the CLL mouse model Eμ-TCL1 decreases leukemic cell invasiveness and disease burden [15,17], suggesting a role for IL-9 in CLL pathogenesis.

The CLL cell secretome, as well as its implication in CTL suppression, is still largely unknown. The characterization of CLL-secreted factors as indirect inhibitors of IS formation might help in developing therapeutic approaches to reactivate anti-tumoral CTL activities in CLL. Here, we hypothesize that soluble factors released by CLL cells mediate CTL disabling. We show that IL-9 secreted by CLL cells from patients with aggressive disease contributes to suppress IS formation and cytotoxic functions of CTLs by enhancing PD-1 expression. Hence, leukemic cell-derived IL-9 concurs to CLL pathogenesis by altering the killing ability of CTLs.

## Results

### Soluble factors released by CLL cells enhance PD-1 expression in CTLs and suppress their ability to assemble functional ISs

CTLs from CLL patients show an exhausted phenotype, with impaired ability to form the IS and kill target cells [3,12]. Accordingly, we observed enhanced expression of the exhaustion markers PD-1, CTLA-4 and LAG-3 in CD8^+^ cells from CLL patients compared to healthy CD8^+^ cells (Supplementary Fig. 1A,B). We asked whether the CLL cell secretome participates in CTL exhaustion. CD8^+^ cells immunopurified from healthy donors were differentiated to CTLs [18] and cultured for 48 h in the presence of complete culture medium alone or media conditioned by either healthy B cells or leukemic cells from CLL patients (experimental workflow in Fig. 1A). Under these experimental conditions healthy CD8^+^ cells acquired the surface expression of the activation markers CD69 and CD25 (Supplementary Fig. 1C) and showed an exhausted phenotype, with enhanced expression of PD-1, CTLA-4 and LAG-3 (Supplementary Fig. 1D-F). Of note, the expression of PD-1 was significantly enhanced in healthy CTLs cultured in CLL cell-conditioned media compared to CTLs cultured in healthy B cell-conditioned media, but not that of CTLA-4 and LAG-3 (Fig. 1B,C, Supplementary Fig. 1D-F), indicating that leukemic cells indirectly suppress CTL function by releasing soluble factors that promote their exhaustion by enhancing PD-1 expression.

**Figure 1.**
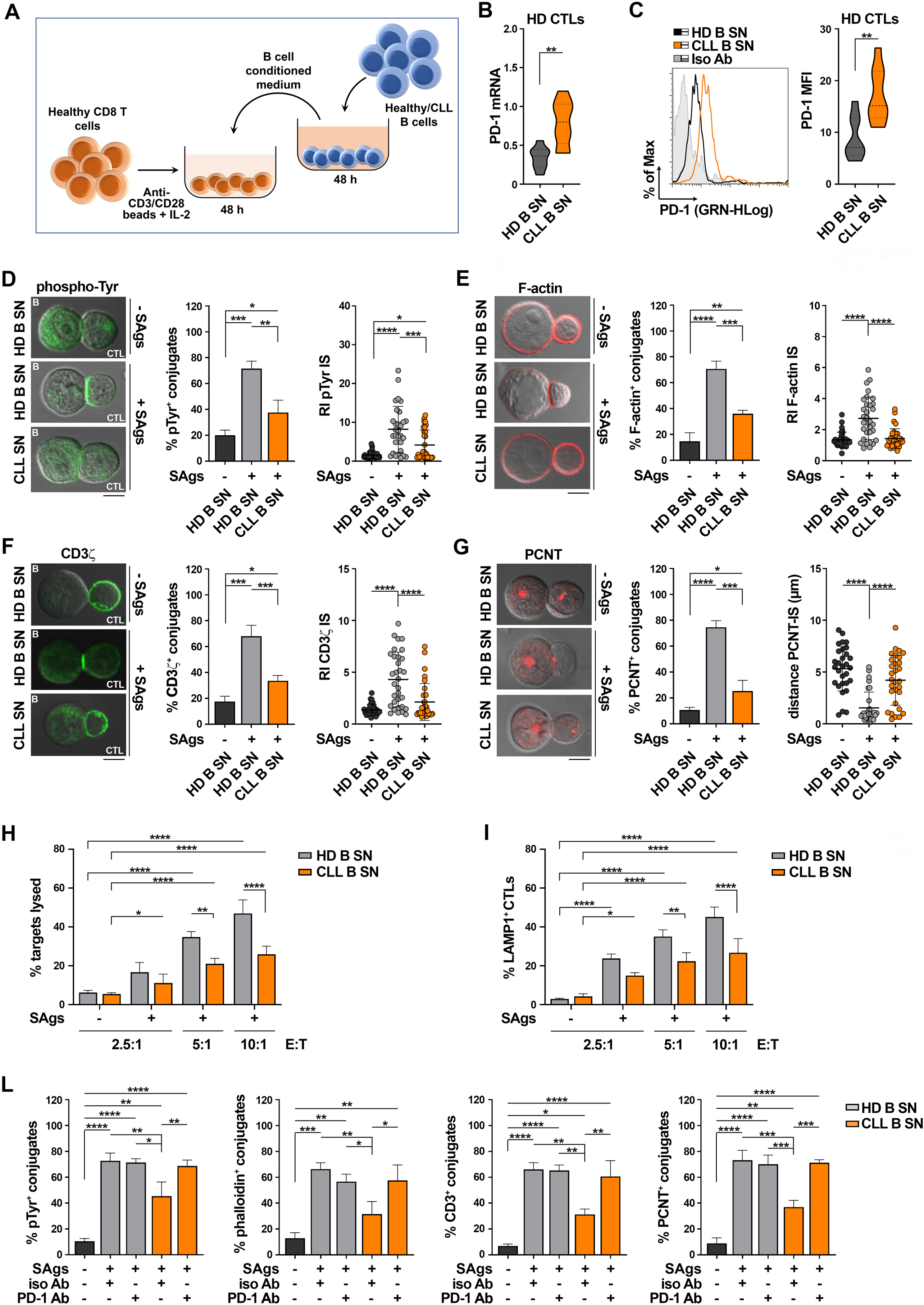
Soluble factors released by CLL cells enhance PD-1 expression on CTLs and suppress their ability to form functional ISs. **A.** Workflow for CTL generation from CD8^+^ cells purified from buffy coats of healthy donors and their treatment with media conditioned by either healthy B cells or B cells purified from CLL patients. **B,C.** qRT-PCR analysis of mRNA (**B**) and flow cytometric analysis of surface (**C**) expression of PD-1 in CD8^+^ cells purified from buffy coats, stimulated with anti-CD3/CD28 mAb-coated beads + IL-2 for 48 h (CTLs), in the presence of conditioned media. A representative flow cytometric histogram is shown (n buffy coats from healthy donors = 9, Mann-Whitney Rank Sum test). **D-G**. Immunofluorescence analysis of pTyr (**D**), F-actin (**E**), CD3ζ. (**F**), and PCNT (**G**) in CTLs activated as in (**A-C**), mixed with Raji cells (APCs) either unpulsed or pulsed with a combination of SEA, SEB, and SEE (SAgs), and incubated for 15 min at 37°C. Data are expressed as % of 15-min SAg-specific conjugates harboring staining at the IS (≥50 cells/sample, n independent experiments = 3, one-way ANOVA test). Representative images (medial optical sections) of the T cell:APC conjugates are shown. Scale bar, 5 μm. **D-F**, *right panels.* Relative fluorescence intensity of pTyr (**D**), F-actin (**E**), and CD3ζ (**F**) at the IS (recruitment index, RI; 10 cells/sample, n independent experiments = 3, one-way ANOVA test). **G**, *right panel*. Measurement of the distance (μm) of the centrosome (PCNT) from the CTL:APC contact in conjugates formed as above (10 cells/sample, n independent experiments = 3, one-way ANOVA test). **H**. Flow cytometric analysis of target cell killing by CTLs cultured for 7 days in conditioned media, using SAg-loaded CFSE-labelled Raji cells as targets at an E:T cell ratio 2.5:1, 5:1 and 10:1. Cells were cocultured for 4 h and stained with propidium iodide prior to processing for flow cytometry. Analyses were carried out gating on CFSE^+^/PI^+^ cells. The histogram shows the percentage (%) of target cells lysed (n≥3, two-way ANOVA test) **I**. Flow cytometric analysis of degranulation of CTLs cultured for 7 days as in (**H**), then cocultured with CFSE-stained SAg-loaded Raji cells for 4 h. The histogram shows the percentage (%) of LAMP1^+^ CTLs, measured gating on the CSFE-negative population (n independent experiments ≥ 3, two-way ANOVA test). **L.** Immunofluorescence analysis of pTyr, F-actin, CD3ζ, and PCNT in CTLs activated for 48 h in the presence of conditioned media, mixed with SAg-loaded Raji cells (APCs), and incubated for 15 min at 37°C. Immediately before mixing the cells, either isotype control (iso Ab) or anti-PD-1 (PD-1 Ab) antibodies were added to CTLs. Data are expressed as % of 15-min SAg-specific conjugates harboring pTyr, F-actin, CD3ζ, and PCNT staining at the IS (≥50 cells/sample, n independent experiments = 3, one-way ANOVA test). Data are expressed as mean±SD. ****, *p* ≤ 0.0001; ***, *p* ≤ 0.001; **, *p* ≤ 0.01; *, *p* ≤ 0.05.

CLL cells suppress the ability of T cells to form functional ISs through cell-to-cell inhibitory contacts [3,4]. The contact-independent enhancement in PD-1 expression in CTLs cultured in conditioned media of CLL cells suggests that soluble CLL-derived factors may also contribute to the IS abnormalities. Healthy CD8^+^ cells were cultured for 48 h in the presence of healthy B cell- or CLL cell-conditioned media and conjugated with Raji cells pulsed with a mixture of the Staphylococcal enterotoxins A, B and E (SAgs). CLL cell-conditioned media significantly impaired IS assembly, as shown by the decreased frequency of conjugates displaying tyrosine phosphoprotein (pTyr), TCR/CD3 and F-actin staining at the CTL/APC interface and the impairment in their synaptic accumulation (Fig. 1D-F). Additionally, centrosome polarization, which is essential for lytic granule delivery to the target cell, was impaired under these conditions, as assessed by measuring the frequency of conjugates displaying centrosome localization beneath the IS membrane and its distance from the IS center (Fig. 1G).

IS suppression was also observed in conjugates of healthy CTLs cultured in vitro in CLL supernatants and allogeneic CLL cells (Supplementary Fig. 2). The IS defects were significantly more pronounced when the conjugation assays were performed using CD8^+^ cells from CLL patients and allogeneic CLL cells (Supplementary Fig. 2), suggesting that in vivo CD8^+^ cell conditioning exacerbates their inability to form the IS, likely through the combination of indirect mechanisms, mediated by CLL-derived soluble factors, and direct interactions involving multiple surface receptor-ligand inhibitory pairs [12].

To test the outcome of these defects on CTL-mediated killing, healthy donor CD8^+^ cells were activated in the presence of healthy or leukemic B cell-conditioned media. After 7 days CTLs were incubated with CFSE-labelled SAg-pulsed Raji cells for 4 h, followed by propidium iodide staining and flow cytometric analysis. Consistent with the IS defects, CLL-conditioned media suppressed the ability of CTLs to kill target cells (Fig. 1H). Moreover, they significantly impaired CTL degranulation, as assessed by measuring the percentage of CD8^+^ cells expressing at their surface the lytic granule marker LAMP-1 (Fig. 1I) [18].

By counteracting TCR-dependent signaling, PD-1 prevents CTLs from forming functional ISs [1,19]. To assess whether PD-1 overexpression induced in healthy CTLs by CLL-conditioned media is causal to their IS assembly defects, PD-1 neutralizing antibodies were added to CTLs cultured as in figure 1A, immediately prior to conjugate formation with SAg-pulsed Raji cells. PD-1 neutralization reverted the IS-suppressive effect of CLL cell-conditioned media (Fig. 1L). Hence, leukemic cells indirectly control CTL exhaustion by secreting soluble factors that enhance PD-1 expression, which in turn impairs the ability of CTLs to form functional ISs and kill target cells.

### The p66Shc expression defect in CLL cells contributes to their soluble PD1-elevating and IS-disrupting activity

CLL cells have a profound defect in the expression of the pro-oxidant adaptor p66Shc, with the lowest levels in leukemic cells from patients with unfavourable prognosis [20]. This defect, which translates into a redox imbalance [21,22], impinges on the expression of a number of CLL-critical genes, including genes encoding lymphoid homing and egress receptors, and cytokines such as IL-9 [17]. Since T cells from poor prognosis patients show higher PD-1 levels compared to favourable prognosis patients [12], we asked whether the p66Shc defect in CLL cells may impact on the expression of the soluble factors that promote PD-1 expression in CTLs.

The mRNA levels of p66Shc in CLL cells inversely correlated with PD-1 mRNA and surface expression in patient-matched CD8^+^ cells (Fig. 2A). This inverse correlation was also observed when healthy CTLs were cultured in media conditioned by CLL cells with different levels of residual p66Shc (Fig. 2B), suggesting the existence of an indirect p66Shc-dependent mechanism through which CLL cells regulate PD-1 expression in CTLs. To test this hypothesis, we reconstituted p66Shc expression in CLL cells by transient nucleofection (Fig. 2C). Media conditioned by CLL transfectants were used to culture healthy CTLs. p66Shc reconstitution reverted the PD-1-enhancing (Fig. 2D,E) and IS-suppressive (Fig. 2F) activities of CLL-conditioned media on CTLs. Hence the p66Shc defect in CLL cells modulates their secretome to promote PD-1 elevation and CTL suppression.

**Figure 2.**
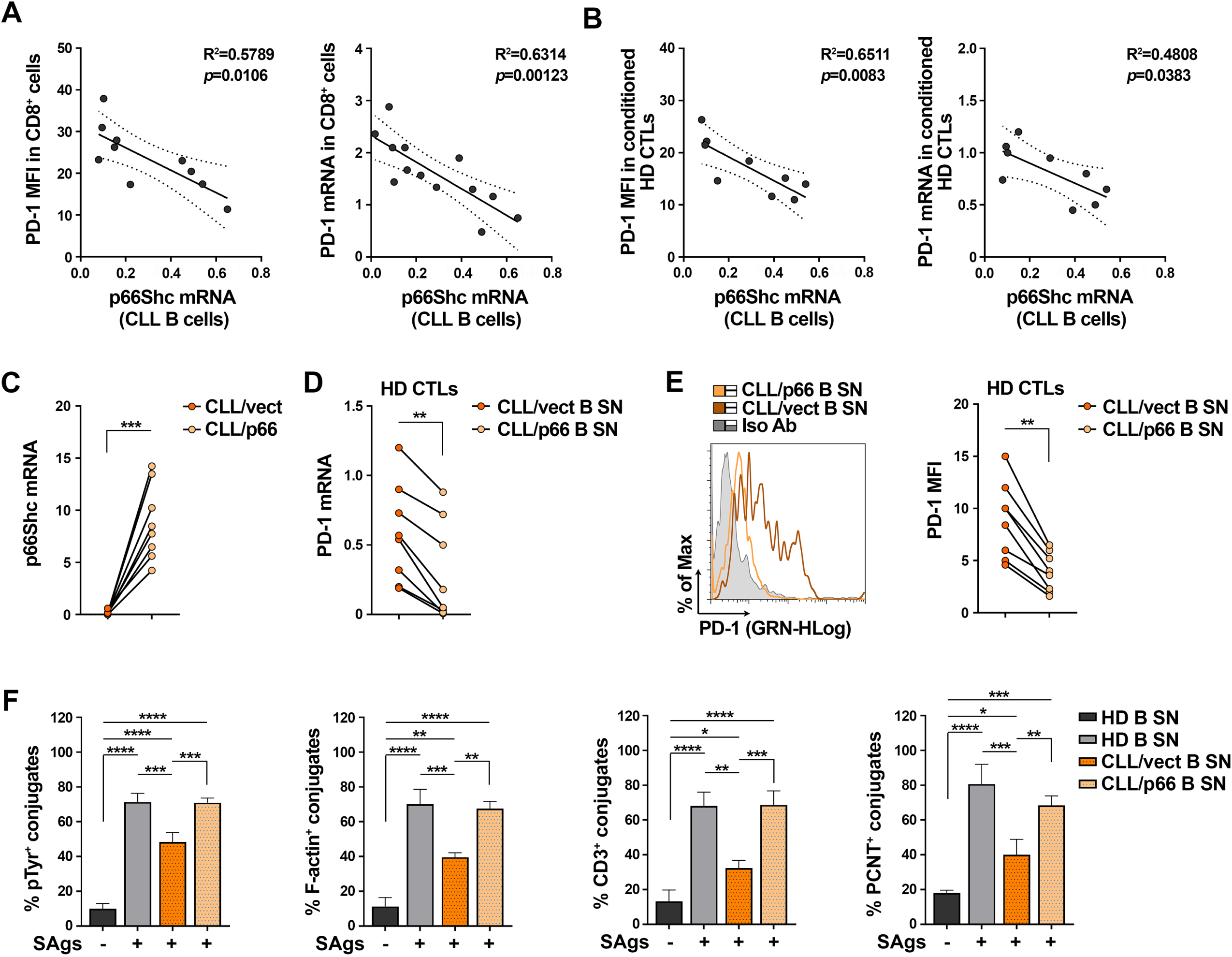
PD-1 overexpression induced in CTLs by CLL culture supernatants is caused by p66Shc deficiency. **A.** Correlation between either surface (MFI) or mRNA levels of PD-1 in CD8^+^ cells isolated from CLL patients and the mRNA levels of p66Shc in B cells purified from the respective CLL patients (n CLL patients ≥ 10). **B**. Correlation between either surface (MFI) or mRNA levels of PD-1 in healthy CTLs activated for 48 h in media conditioned by CLL-B cells, and the mRNA levels of p66Shc in B cells purified from the respective CLL patients used to generate conditioned media (n=9). **C.** qRT-PCR analysis of p66Shc mRNA in B cells purified from CLL patients (n=8) and transfected with either empty (CLL/vect) or p66Shc-encoding (CLL/p66) vectors (paired t test). **D, E.** qRT-PCR analysis of mRNA (**E**) and flow cytometric analysis of surface (**F**) expression of PD-1 in CTLs from heathy donors (n=8), activated for 48 h in the presence of media conditioned by CLL B cell transfectants. Representative flow cytometric histograms are shown (paired t test). **F.** Immunofluorescence analysis of pTyr, F-actin, CD3ζ, and PCNT in CTLs activated in media conditioned by CLL-B cell transfectants, mixed with SAg-loaded Raji cells (APCs), and incubated for 15 min at 37°C. Data are expressed as % of 15-min SAg-specific conjugates harboring pTyr, F-actin, CD3ζ, and PCNT staining at the IS (≥ 50 cells/sample, n independent experiments=3, one-way ANOVA test). Data are expressed as mean±SD. ****, *p* ≤ 0.0001; ***, *p* ≤ 0.001; **, *p* ≤ 0.01; *, *p* ≤ 0.05.

### p66Shc deficiency in leukemic cells from Eμ-TCL1 mice enhances their soluble PD1-elevating activity on CD8^+^ cells

In agreement with a previous report [23], we found PD-1 overexpressed in CD8^+^ cells isolated from lymph nodes of Eμ-TCL1 mice, the mouse model of CLL [24], compared to their wild-type counterparts (Fig 3A,B). PD-1 expression increased during disease progression, as assessed by flow cytometric analysis of CD8^+^ cells from Eμ-TCL1 mice with either mild (20-39% leukemic cells in PB, 1-2×10^7^ WBC/ml PB) or overt (>40% leukemic cells in PB, >2×10^7^ WBC/ml PB) leukemia (Fig. 3C, Supplementary Fig. 3A). PD-1 mRNA and surface expression directly correlated with the percentage of circulating leukemic cells (Fig. 3D).

**Figure 3.**
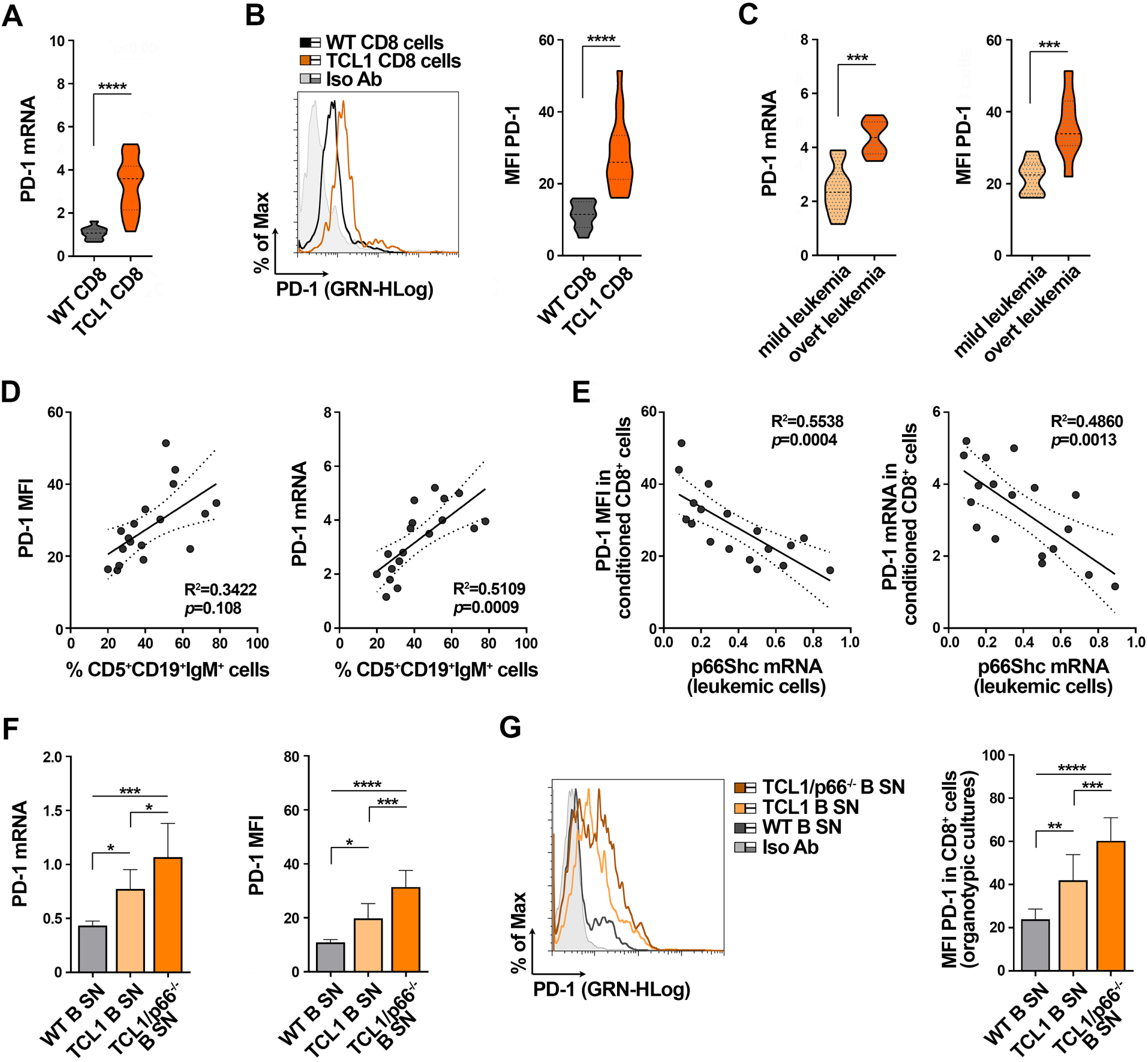
p66Shc deficiency in Eμ-TCL1 mice enhances the ability of leukemic cells to enhance PD-1 expression in CD8^+^ cells. **A,B.** qRT-PCR analysis of mRNA (**A**) and flow cytometric analysis of surface (**B**) expression of PD-1 in CD8^+^ cells isolated from lymph nodes of either wild-type (n = 10) or Eμ-TCL1 (n = 18) mice. Representative flow cytometric histograms are shown (Mann Whitney Rank Sum test). **C**. mRNA expression of PD-1 in CD8^+^ cells from lymph nodes of Eμ-TCL1 shown in (**A)** were subgrouped, according to disease stage, in mild leukemia (20-39% leukemic cells in PB, 1-2×10^7^ WBC/ml PB, n Eμ-TCL1 mice = 10) and overt leukemia (>40% leukemic cells in PB, >2×10^7^ WBC/ml PB, n Eμ-TCL1 mice = 8). (Mann Whitney Rank Sum test). **D**. Correlation between surface (MFI) or mRNA levels of PD-1 in CD8^+^ cells isolated from lymph nodes of Eμ-TCL1 mice and the percentage of leukemic cells in peripheral blood of the same mouse. **E**. Correlation between surface (MFI) or mRNA levels of PD-1 in CD8^+^ cells isolated from lymph nodes of wild-type mice cultured in the presence of media conditioned by splenic B cells from Eμ-TCL1 mice, and the mRNA levels p66Shc in Eμ-TCL1 cells used to generate conditioned media (n wild-type mice=18). **F**. qRT-PCR analysis of PD-1 mRNA in CD8^+^ cells isolated from spleens of wild-type mice (n=10), cultured in the presence of media conditioned by splenic B cells from either wild-type, Eμ-TCL1 or Eμ-TCL1/p66^-/-^ mice (n independent experiments ≥5; one-way ANOVA). **G**. Flow cytometric analysis of PD-1^+^CD8^+^ cells in 220 μm-thick slices from spleens of wild-type mice (n=10), cultured as in (**F**) (n independent experiments ≥5; one-way ANOVA). Data are expressed as mean±SD. ****, *p* ≤ 0.0001; ***, *p* ≤ 0.001; **, *p* ≤ 0.01; *, *p* ≤ 0.05.

We previously reported that a p66Shc expression defect in leukemic cells from Eμ-TCL1 mice, that becomes more pronounced during disease progression (Supplementary Fig. 3B) [25], contributes to disease development and aggressiveness [17,20,25]. Consistent with the data obtained in CLL cells, PD-1 mRNA and surface expression in CD8^+^ cells inversely correlated with the levels of residual p66Shc mRNA in leukemic Eμ-TCL1 cells (Fig. 3E).

We asked whether the suppressive activity of CLL cell-conditioned media was reproduced in Eμ-TCL1 mice. PD-1 expression was enhanced in lymph node CD8^+^ cells purified from wild-type mice cultured in Eμ-TCL1 cell-conditioned media (Fig. 3F) (same experimental workflow as for human CD8^+^ cells, Fig. 1A), indicating that, similar to human CLL, leukemic cells promote bystander CD8^+^ cell exhaustion in Eμ-TCL1 mice through a contact-independent mechanism. In support of this notion, adoptively transferred CD8^+^ cells purified from spleens of OT-1 mice that express a CLL-irrelevant TCR in our model showed enhanced PD-1 expression in Eμ-TCL1 mice with overt leukemia compared to wild-type mice (Supplementary Fig. 4).

PD-1 expression further increased in healthy splenic CD8^+^ cells cultured in media conditioned by leukemic cells from Eμ-TCL1/p66Shc^-/-^ mice (Fig. 3F), which develop an aggressive disease [25]. Similar results were obtained using organotypic cultures from spleens of wild-type mice, a system that recapitulates the key physiological features of lymphoid organs [26], cultured in media conditioned by B cells from wild-type, Eμ-TCL1 or Eμ-TCL1/p66Shc^-/-^ mice (Fig. 3G). Hence, leukemic cells from Eμ-TCL1 mice indirectly suppress CD8^+^ cell functions by secreting soluble factors that enhance PD-1 expression in a p66Shc-dependent manner.

### p66Shc deficiency impinges on the cytokine landscape of CLL cells

CLL cell-derived IL-9 promotes the secretion of homing chemokines by stromal cells of the TME [17]. To identify leukemic cell-derived soluble factors with potential CTL-suppressive activity, we first quantified a panel of cytokines and chemokines released in media conditioned by splenic B cells from wild-type, Eμ-TCL1 or Eμ-TCL1/p66^-/-^ mice by Multiplex ELISA. We identified soluble factors whose amounts were significantly increased in media conditioned by leukemic cells compared to their wild-type counterparts (Fig. 4A, Supplementary Table 1), and selected the top candidates based on their significantly higher amounts in Eμ-TCL1 cell- *vs* wild-type B cell-conditioned media, and Eμ-TCL1/p66^-/-^ cell- *vs* Eμ-TCL1 cell-conditioned media. Moreover, we established a threshold of 100 ng/ml in media conditioned by wild-type cells. Among all soluble factors tested, only CCL22, CCL24, IL-9 and IL-10 met all our criteria. Of these, CCL22 and IL-9 were previously shown to be overexpressed by leukemic cells isolated from spleens of Eμ-TCL1/p66Shc^-/-^ compared to Eμ-TCL1 mice (Fig. 4B, Supplementary Table 2 and [17]).

**Figure 4.**
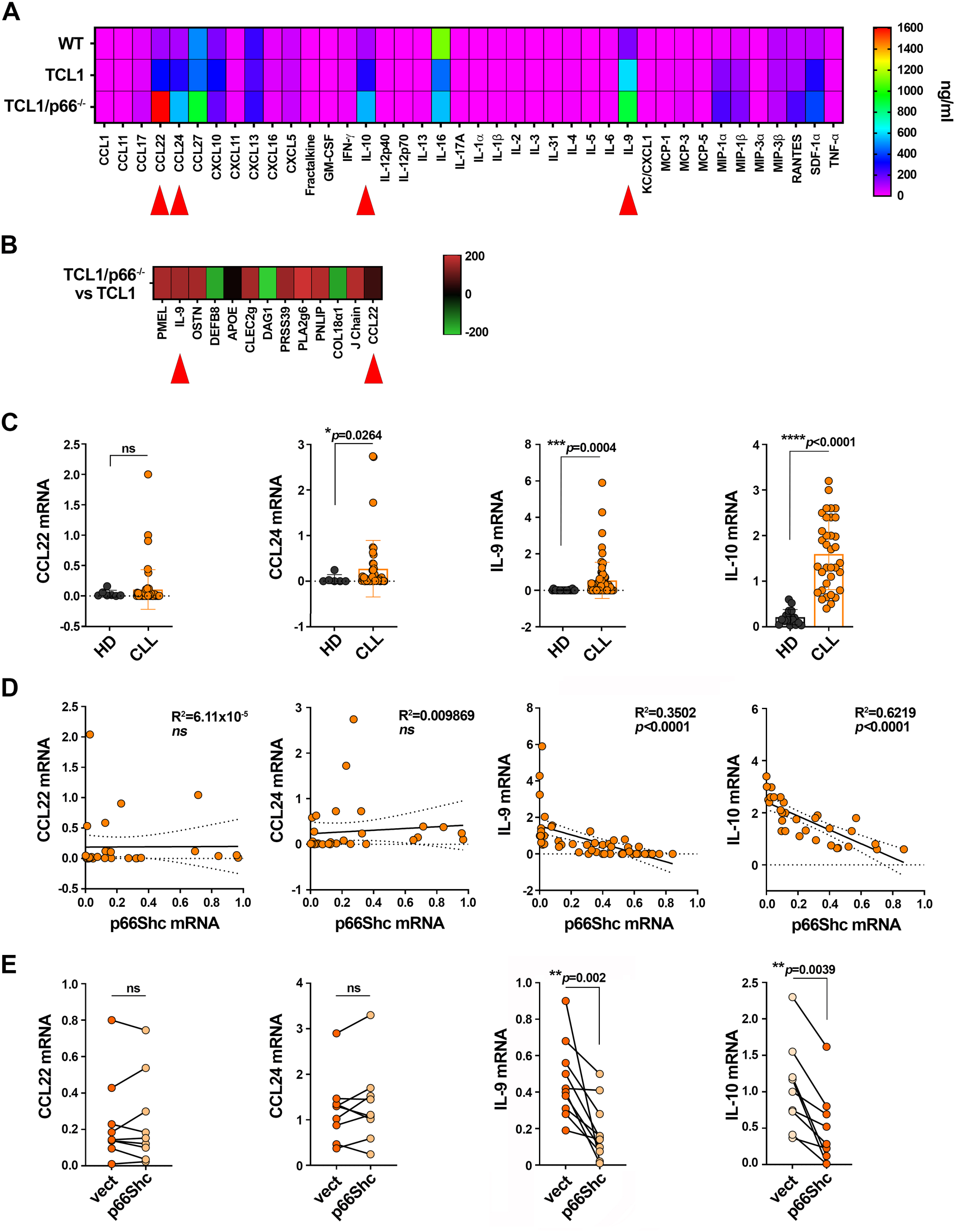
p66Shc deficiency impinges on the cytokine landscape of leukemic cells in Eμ-TCL1 mice and CLL. **A.** Heat map of the amounts of cytokines and chemokines in the culture supernatants of B cells isolated from spleens of wild-type mice (WT; n=40), or leukemic cells purified from spleens of Eμ-TCL1 (TCL1; n=40) or Eμ-TCL1/p66^-/-^ (TCL1/p66^-/-^; n=40) mice and quantified by Multiplex ELISA. **B**. Heat map of transcripts from Affymetrix array analysis showing differential expression patterns between leukemic Eμ-TCL1 (TCL1, n=3) and Eμ-TCL1/p66Shc^-/-^ (TCL1/p66^-/-^, n=3) cells. Differential expression criteria: *p*-value < 0.05, estimated fold change > 2. Up-regulated and down-regulated transcripts are shown in red and green, respectively. **C**. qRT-PCR analysis of mRNA expression of CCL22, CCL24, IL-9 and IL-10 in B cells purified from peripheral blood of healthy donors (n≥6) or CLL patients (n≥45) (Mann Whitney Rak Sum test). **D.** Correlation between mRNA levels of CCL22, CCL24, IL-9 or IL-10 and p66Shc mRNA levels in the respective CLL patients (n≥31). **E**. qRT-PCR analysis of mRNA expression of CCL22, CCL24, IL-9 and IL-10 in B cells purified from CLL patients (n=9) and transiently nucleofected with either empty (vect) or with p66Shc-expressing (p66Shc) vectors (Wilcoxon test). Data are expressed as mean±SD. ****, *p* ≤ 0.0001; ***, *p* ≤ 0.001; **, *p* ≤ 0.01; *, *p* ≤ 0.05.

To translate these findings to the context of CLL, we quantified the selected candidates in healthy and CLL cells by qRT-PCR. The levels of CCL24, IL-9 and IL-10 mRNAs were significantly higher in CLL cells compared to healthy B cells, while CCL22 expression was comparable to healthy B cells (Fig. 4C). As previously reported [17], the IL-9 mRNA levels inversely correlated with the p66Shc mRNA levels (Fig. 4D). This also applied to IL-10, but not to CCL22 or CCL24 (Fig. 4D). Moreover, p66Shc reconstitution in CLL cells by transient nucleofection led to a decrease in the mRNA of IL-9 (Fig. 4E and [17]) and IL-10, but not of CCL22 or CCL24 (Fig. 4E).

### IL-9 secreted by leukemic cells from CLL patients promotes PD-1 expression in CTLs

Based on these results, we assessed IL-9 and IL-10 as potential p66Shc-dependent, CTL-suppressive soluble factors produced by CLL cells. We quantified surface PD-1 in healthy CTLs cultured with healthy or leukemic B cell-conditioned media, in the presence of anti-IL-9 or anti-IL-10 neutralizing antibodies. Anti-IL-9 (Fig. 5A) but not anti-IL-10 (Fig. 5B) antibodies impaired the PD-1-elevating activity of CLL cell supernatants in healthy CTLs. These data were confirmed using recombinant IL-9 that, as opposed to recombinant IL-10, enhanced surface PD-1 expression in healthy CTLs cultured in healthy B cell-conditioned media (Fig. 5A,B). These data indicate that IL-9, but not IL-10, mediates the PD-1- elevating ability of leukemic cells on healthy CTLs.

**Figure 5.**
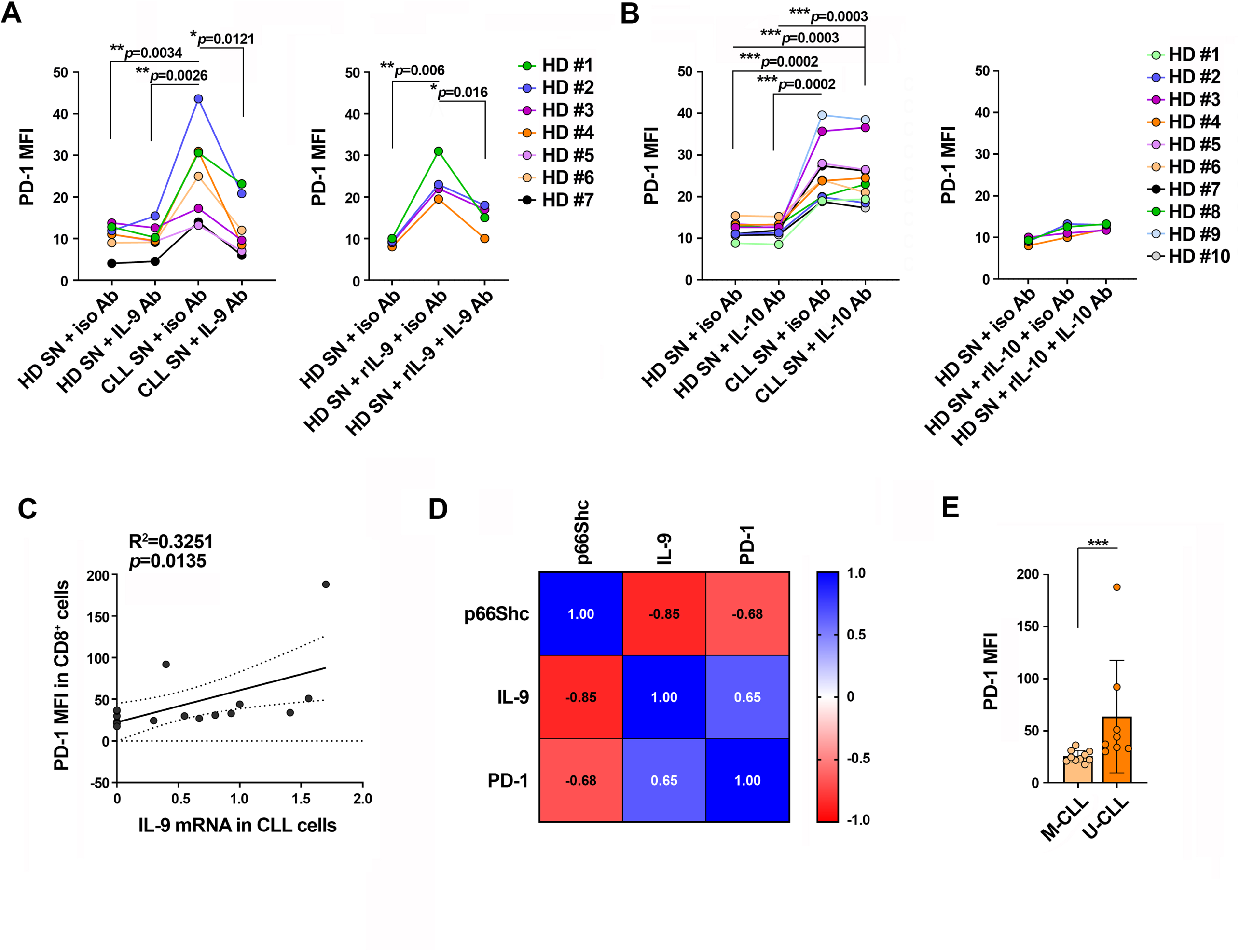
IL-9 secreted by CLL cells promotes PD-1 expression in CTLs. **A,B**. Flow cytometric analysis of surface PD-1 in CD8^+^ cells purified from buffy coats, stimulated with anti-CD3/CD28 mAb-coated beads + IL-2 for 48 h (CTLs), and cultured in conditioned media in the presence of either control (iso Ab), anti-IL-9 (IL-9 Ab, **A**) or anti-IL-10 (IL-10 Ab, **B**) antibodies and/or recombinant IL-9 (rIL-9) (**A**), or recombinant IL-10 (rIL-10) (**B**) (n buffy coats from healthy donors = 4; one-way ANOVA). **C**. Correlation between surface PD-1 (MFI) in CD8^+^ cells isolated from CLL patients and IL-9 mRNA in leukemic cells isolated from the respective CLL patients (n=18). **D**. Spearman r correlation index between surface PD-1 (MFI) in CD8^+^ cells isolated from CLL patients and IL-9 and p66Shc mRNA in CLL cells from the respective CLL patients (n=18). **E**. Flow cytometric analysis of surface PD-1 in CD8^+^ cells isolated from peripheral blood of CLL patients, grouped in M-CLL (n=10) and U-CLL (n=8) patients according to the *IGHV* mutational status. Data are expressed as mean±SD. ****, *p* ≤ 0.0001; ***, *p* ≤ 0.001; **, *p* ≤ 0.01; *, *p* ≤ 0.05.

We correlated PD-1 expression on CD8^+^ cells with IL-9 expression in CLL cells. PD-1 expression on CD8^+^ cells directly correlated with IL-9 expression in patient-matched leukemic cells (Fig. 5C, Supplementary Table 3). The heat map shown in figure 4G, which correlates PD-1 expression in CD8^+^ cells with both IL-9 and p66Shc expression in patient-matched leukemic cells, highlighted a strong correlation among the three markers, with PD-1 expression on CD8^+^ cells directly correlated with IL-9 but inversely correlated with p66Shc expression in leukemic cells (Fig. 5D, Table 3). PD-1 expression was higher in CD8^+^ cells from patients with unmutated *IGHV* genes (U-CLL), a marker of early relapse after fixed-duration therapy (Fig.5E), and express the lowest p66Shc levels, compared to patients with mutated *IGHV* genes (M-CLL) (Table 3) [20,27]. These results highlight a new mechanism exploited by CLL cells to disable the tumor-suppressive activity of CTLs involving IL-9 secretion into the TME to enhance PD-1 expression in CTLs.

### IL-9 secreted by leukemic cells from CLL patients impairs IS formation in CTLs

CLL cells suppress the ability of activated CD8^+^ cells to assemble the IS through direct interaction of surface inhibitory receptor/ligand axes (Fig. 1L) [12]. To test the hypothesis that leukemic cell-derived IL-9 contributes to the assembly of dysfunctional ISs by modulating PD-1 expression in CTLs, healthy CD8^+^ cells were cultured with healthy B cell or CLL cell-conditioned media in the presence of either isotype control or anti-IL-9 neutralizing antibodies prior to conjugate formation. Anti-IL-9 antibodies counteracted the suppressive activities of leukemic cell supernatants on IS formation (Fig. 6A-D, left panels). Consistent with these findings, recombinant IL-9 added to healthy B cell-conditioned media impaired IS formation, an effect that was not observed in the presence of neutralizing anti-IL-9 antibodies (Fig. 6A-D), Consistent with the IS defects (Fig. 6A-D), the suppressive activity of the CLL supernatants on CTL-mediated killing and degranulation were neutralized by anti-IL-9 antibodies (Fig. 6E-F). Moreover, recombinant IL-9 reproduced the suppressive activities of CLL supernatants (Fig. 6E-F). Collectively, these results demonstrate that CLL cell-derived IL-9 impairs IS formation and effector functions of CTLs by promoting PD-1 expression in CTLs.

**Figure 6.**
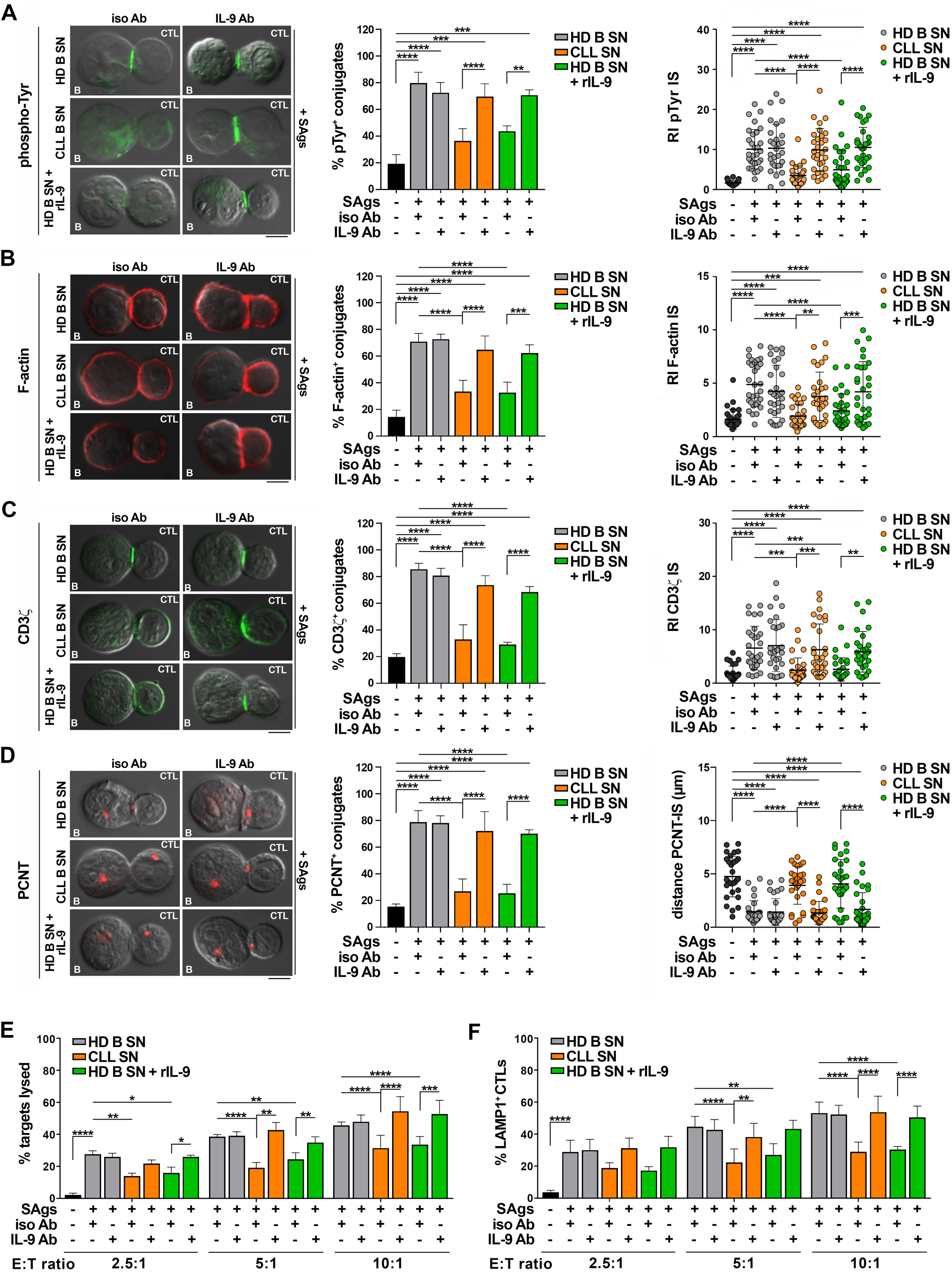
IL-9 secreted by CLL cells suppresses IS formation and effector functions in CTLs. **A-D,** *left panels.* Immunofluorescence analysis of pTyr (**A**), F-actin (**B**), CD3ζ(**C**), and PCNT (**D**) in CTLs activated for 48 h in the presence of conditioned media, with the addition of either control (iso Ab), anti-IL-9 (IL-9 Ab) or recombinant IL-9 (rIL-9), mixed with Raji cells (APCs) either unpulsed or pulsed with a combination of SAgs, and incubated for 15 min at 37°C. Data are expressed as % of 15-min SAg-specific conjugates harboring staining at the IS (≥50 cells/sample, n independent experiments ≥3, one-way ANOVA test). Representative images (medial optical sections) of the T cell:APC conjugates are shown. Scale bar, 5 μm. **A-C**, *right panels.* Relative fluorescence intensity of pTyr (**A**), F-actin (**B**), and CD3ζ (**C**) at the IS (recruitment index, RI; 10 cells/sample, n independent experiments ≥3, one-way ANOVA test). **D**, *right panel*. Measurement of the distance (μm) of the centrosome (PCNT) from the CTL:APC contact in conjugates formed as above (10 cells/sample, n independent experiments =3, one-way ANOVA test). **E**. Flow cytometric analysis of target cell killing by CTLs cultured for 7 days in conditioned media, with the addition of either control (iso Ab), anti-IL-9 (IL-9 Ab) or recombinant IL-9 (rIL-9), using SAg-loaded Raji cells as targets at an E:T cell ratio 2.5:1, 5:1 and 10:1. Cells were cocultured for 4 h and stained with propidium iodide prior to processing for flow cytometry. Analyses were carried out gating on CFSE^+^PI^+^ cells. The histogram shows the percentage (%) of target cells lysed (n independent experiments =5, two-way ANOVA test). **F**. Flow cytometric analysis of degranulation of CTLs cultured for 7 days as in (**E**), then cocultured with CFSE-stained Raji cells loaded with SAg at an E:T cell ratio 2.5:1, 5:1 and 10:1 for 4 h. The histogram shows the percentage (%) of LAMP1^+^ CTLs, measured gating on the CSFE-negative population (n=6, two-way ANOVA test). Data are expressed as mean±SD. ****, *p* ≤ 0.0001; ***, *p* ≤ 0.001; **, *p* ≤ 0.01; *, *p* ≤ 0.05.

### In vivo IL-9 blockade in Eμ-TCL1/p66^-/-^ mice normalizes PD-1 expression in CD8^+^ cells

To validate the role of IL-9 in PD-1 overexpression in CD8^+^ cells in vivo, we carried out IL-9-blockade experiments in the aggressive CLL Eμ-TCL1/p66Shc^-/-^ mouse model [28], [17]. Mice with overt disease were intraperitoneally administered anti-IL-9 or isotype-control mAbs twice a week for 4 weeks (Fig. 7A) [17]. Flow cytometry analysis of PD-1^+^CD8^+^ splenocytes demonstrated that IL-9 blockade resulted in a reduction in PD1 expression and frequency of PD-1^+^CD8^+^ cells compared to isotype control (Fig. 7B). Enhanced surface PD-1 expression in CD8^+^ cells was also observed in in vitro experiments where organotypic spleen cultures from wild-type mice were cultured for 48 h in media conditioned by leukemic cells from Eμ-TCL1/p66Shc^-/-^ mice (Fig. 7C), that were enriched in IL-9 (Fig. 7D). A similar PD-1 enhancement was obtained when wild-type spleen slices were cultured in media conditioned by wild-type B cells, where IL-9 is barely detectable (Fig. 7D), in the presence of recombinant IL-9 (Fig. 7C). Anti-IL-9 antibodies neutralized the enhancement in PD-1 expression induced both by Eμ-TCL1/p66Shc^-/-^ leukemic cell supernatants and by wild-type B cell supernatants added with IL-9 (Fig. 7C), indicating that leukemic cell-derived, IL-9-rich supernatants can shape the splenic microenvironment to a pro-tumoral one.

**Figure 7.**
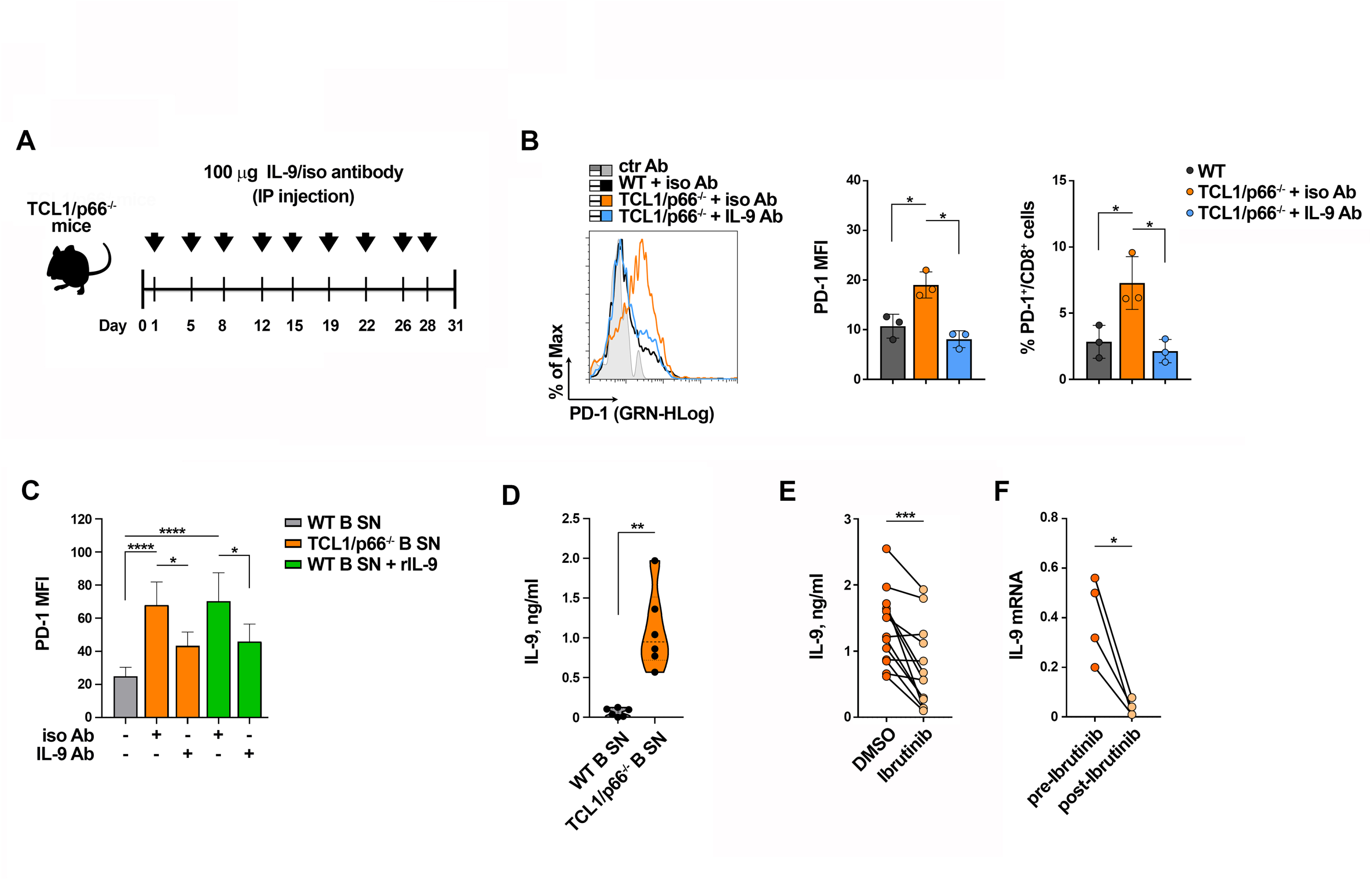
In vivo IL-9 blockade in Eμ-TCL1/p66^-/-^ mice normalizes PD-1 expression in CD8^+^ cells. **A.** Schematic representation of the experimental design of intraperitoneal injection of wild-type C57BL/6 mice (n=3) or Eμ-TCL1/p66Shc^-/-^ mice (n=6) with overt leukemia (∼40% CD5^+^CD19^+^ leukemic cells in PB) with 100 μg of either isotype control (n=3) or anti-IL-9 (n=3) antibodies dissolved in 100 μl PBS. **B**. Flow cytometric analysis of PD-1^+^CD8^+^ cells in spleens isolated from either wild type mice injected with PBS (n=3) or Eμ-TCL1/p66Shc^-/-^ mice (n=6) injected with anti-IL-9 (IL-9 Ab, n=3) or isotype (iso Ab, n=3) mAb. Data are expressed as MFI of PD-1 in CD8^+^-gated cells (left) and as % PD-1^+^CD8^+^ cells (right) (RM one-way ANOVA). **C**. Flow cytometric analysis of PD-1^+^CD8^+^ cells in 220 μm-thick slices obtained from spleens of wild-type mice (n=3) and cultured for 48 h in the presence of conditioned media, with the addition of either control (iso Ab), anti-IL-9 (IL-9 Ab) or recombinant IL-9 (rIL-9) (n independent experiments ≥3; one-way ANOVA). **D**. Quantification by ELISA of IL-9 released in media conditioned by splenic B cells from either wild-type (n=6) or Eμ-TCL1/p66^-/-^ (n=6) mice. Mann Whitney Rank Sum test. **E**. Quantification by ELISA of IL-9 released in the culture supernatants of leukemic cells isolated from peripheral blood of CLL patients (n=13) and treated in vitro for 48 h with either DMSO or 10 μM Ibrutinib. Paired t test. **F**. qRT-PCR analysis of mRNA expression of IL-9 in leukemic cells isolated from peripheral blood of CLL patients (n=4) who received 420 mg Ibrutinib once a day. Blood samples were collected on the starting day of Ibrutinib treatment (before Ibrutinib) and at follow-up of Ibrutinib treatment (follow-up Ibrutinib; range follow-up 26.2±7.3 months). Paired t test. Data are expressed as mean±SD. ****, *p* ≤ 0.0001; **, *p* ≤ 0.01; *, *p* ≤ 0.05.

### Ex vivo and in vivo inhibition of BTK enhances IL-9 expression in leukemic cells from CLL patients

Treatment of CLL patients with BTK inhibitors relieves T cell exhaustion by downregulating PD-1 expression [29,30]. We hypothesized that this could partly or completely account for the enhanced release of IL-9 in the TME by leukemic cells. To test this hypothesis, IL-9 secreted in media conditioned by CLL cells treated in vitro for 48 h with the BTK inhibitor Ibrutinib was quantified by ELISA. Ibrutinib treatment decreased IL-9 release by CLL cells (Fig. 7E). Moreover, IL-9 expression was reduced in 4 CLL patients showing a significant response to second line Ibrutinib treatment (range follow-up 26.2±7.3 months) (Fig. 7F, Supplementary Table 4), suggesting that the response of CLL patients to BTK inhibitors results, at least in part, from the normalization of IL-9 release by leukemic cells.

## Discussion

During their growth, tumors develop mechanisms for harnessing immune responses. CTLs are subjected to these suppressive actions, as demonstrated by impairment in their anti-tumoral functions [1]. Direct inhibitory receptor/ligand axes contribute to CTL suppression by inhibiting TCR-dependent signaling pathways [1]. Here we identify a new cell contact-independent mechanism of CLL cell escape from CTL-mediated killing.

Abnormal IL-9 secretion by leukemic cells of CLL patients [17,31–33] and Eμ-TCL1 mice [17] correlates with aggressive disease hallmarks, such as unmutated *IGHV*, low p66Shc levels [15,17], and lower overall survival [17]. The pro-tumoral activity of IL-9 involves its ability to promote homing chemokine secretion by stromal cells of lymphoid organs [15,17], thereby attracting CLL cells to the pro-survival lymphoid niche. Here we found that IL-9 secreted by CLL cells also participates in their escape from CTL-mediated killing by hampering IS formation in CTLs. Abnormal serum IL-9 levels in CLL patients are also associated to uncontrolled secretion by Th-9 cells [33], suggesting that CLL and Th-9 cells potentially cooperate to shape the TME by suppressing CTLs.

CLL cells secrete both anti-tumoral and pro-tumoral cytokines [13]. IL-15 has a frank anti-tumoral activity, as witnessed by enhanced in vitro antibody-dependent cytotoxicity against CLL cells [34]. Moreover, low IL-27 serum levels in CLL patients have been related to the development of aggressive disease and decreased anti-leukemic cytotoxicity of CD8^+^ cells [35]. Conversely, increased IL-6 in CLL sera correlates with a repressed functional profile of circulating CTLs [36,37]. Whether IL-6 participates directly in CTL suppression in CLL has not been assessed to date. Of note, we show that Eμ-TCL1 and wild-type B cells secrete comparable and very low IL-6 amounts, ruling out leukemic cells as the source of IL-6 in CLL sera. On the other hand, our data highlight IL-9, which is overexpressed in CLL patients with aggressive disease presentation [17,31,33], as a key pro-tumoral cytokine that shapes the TME not only to promote leukemic cell accumulation and survival [17], but also to help them evade elimination by CTLs.

PD-1 is almost absent in resting T cells while it is a hallmark of effector T cells [1,38,39]. Several stimuli upregulate PD-1 expression, the most important being TCR-mediated signaling, which activates transcription factors implicated in PD-1 expression, including nuclear factor of activated T cells 1 (NFATC1) [40,41], NF-κB [42], and T-bet [41]. Cytokines such as IL-10 [43] and TGF-β [43,44] have been also found to promote PD-1 expression. Here we show that IL-9 secreted by leukemic cells from CLL patients with aggressive disease presentation enhances PD-1 expression in healthy CTLs, thereby contributing to shift them to the exhausted and disabled phenotype. Our results suggest that agents targeting IL-9, such as anti-IL-9 neutralizing antibodies, are plausible partners for combination therapies with immune checkpoint inhibitors to reverse the state of exhaustion and reinvigorate protective immune responses in CLL. In support of this notion, suppression of IL-10 expression has been shown to enhance anti-tumor T cell immunity in combination with PD-L1 blockade in a preclinical mouse model [45]. Of note, while in vivo IL-10 suppression alone increased the frequency of CD8^+^ cells displaying effector markers, it did not affect PD-1 upregulation [45], consistent with our findings.

Despite the accumulation of PD1^+^CD8^+^ cells in peripheral lymphoid organs of CLL patients, PD-1/PD-L1 blockade alone has shown limited clinical efficacy, with the exception of the most aggressive disease development, Richter’s transformation [46], suggesting that the neutralization of the PD-1 inhibitory signaling axis may not be sufficient to reactivate the CTL anti-tumoral functions, and pointing out the relevant implication of exhaustion molecules other than PD-1, such as CTLA-4 and LAG-3 [8,47,48], in restraining CTL functions. Our results showing a recovery of the ability of isolated exhausted CTLs to form functional ISs in the presence of neutralizing anti-PD-1 antibodies underscore the crucial role of PD-1 in the suppression of anti-tumor CTL activities in the complex context of the TME. Moreover, CTLA-4 and LAG-3 expression was not further increased by conditioned media from CLL cells, ruling them out as mediators of soluble factor-mediated CTL exhaustion in CLL. Nevertheless, a role for other inhibitory receptor/ligand axes, including both membrane-bound and soluble immune checkpoint molecules such as sCTLA-4 (soluble CTLA-4) and sPD-1 (soluble PD-1) [49], cannot be excluded.

The finding that the p66Shc expression defect in CLL cells impinges on their ability to suppress CTLs further witnesses to the key role of this adaptor in leukemic cell shaping of the TME to their advantage. p66Shc is a molecular adaptor that promotes apoptosis through an adaptor-independent pro-oxidant activity [21,22], which contributes to the intrinsic apoptosis defects and chemoresistance of CLL cells [17,27,28]. The ROS-elevating, gene-modulating activity of p66Shc also indirectly favours leukemic cell survival by setting an imbalance between the homing and egress receptors that coordinate lymphocyte trafficking, promoting their accumulation in the pro-survival lymphoid niche [50,51]. Here we show that *p66Shc* deletion in Eμ-TCL1 mice also affects the expression of CLL-critical cytokines such as IL-9 and IL-10 by leukemic cells, which also applies to the p66Shc-deficient CLL cells. This highlights the ramifications in the outcome of the p66Shc defect in CLL cells on strategic processes that participate in their survival.

In summary, our data highlight a crosstalk between leukemic cells and CTLs in CLL that indirectly suppresses the killing activity of CTLs, supporting leukemic cell escape from immunosurveillance and thus contributing to CLL pathogenesis.

## Methods

### CLL patients, healthy donors, mice and cell lines

CLL diagnosis and mutational *IGHV* status were assessed as reported [25,52,53]. PB samples were collected from 96 treatment-naive CLL patients (>95% leukemic CD5^+^CD19^+^/CD19^+^ cells) and 4 CLL patients subjected to Ibrutinib treatment (Supplementary Table 4). Healthy control B cells were purified from 119 buffy coats as described [54]. Transfections were previously described [50,53]. Non-randomized non-blinded experiments were carried out on cells isolated from Eμ-TCL1, Eμ-TCL1/p66Shc^-/-^ mice [24,53], OT-1 mice [55,56] and parental C57BL/6J mice. Disease development and overt leukemia achievement were assessed as reported [25]. In vivo treatment of Eμ-TCL1/p66Shc^-/-^ leukemic mice with anti-IL-9 or isotype ctr antibodies was reported [17]. The Raji B lymphoblastoid cell line was previously described [57].

### Conditioned supernatants and multiplex assays

Conditioned supernatants were generated and stored as reported [17]. Viability of healthy B cells and CLL cells used to generate conditioned supernatants was consistently >65% at 48 h versus ∼85% immediately after purification (Supplementary Fig. 5). Mouse cytokines and chemokines were quantified by Bio-Plex Chemokine Panel Multiplex assay (BioRad). Data were acquired and analyzed using Luminex-Magpix (BioRad).

### Purification, activation and conditioning of CD8^+^ cells

CD8^+^ cells isolated from mouse lymph nodes by immunomagnetic sorting were incubated for 48 h in conditioned supernatants. 0.5 ng/ml recombinant IL-9 and 0.1 ng/ml control or anti-IL-9 mAbs (Supplementary Table 5) were added to culture media. CD8^+^ cells isolated from PB of healthy donors were stimulated with anti-CD3/CD28 beads and 50 U/ml IL-2 and cultured in conditioned media. 20 ng/ml recombinant IL-9, 0.5 ng/ml recombinant IL-10, 0.1 ng/ml isotype control, or 1 ng/ml anti-IL-9 or anti-IL10 mAbs, were added to culture media. 48 h after activation, beads were removed and CTLs collected. For cytotoxicity and degranulation assays, CTLs were expanded for additional 3 days with IL-2, and collected at day 7.

### IS formation and immunofluorescence

Conjugates between CTLs and superantigen (SAg)-pulsed Raji B cells were carried out as detailed in Supplementary Methods [18]. Briefly, Raji cells were loaded with SAgs, mixed with CTLs (1:1.5 APC:CTL ratio) and incubated for 15 min at 37°C. Anti-PD1 mAbs were added during conjugate formation. Samples were seeded onto poly-L-lysine-coated slides, fixed, permeabilized, stained with antibodies (listed in Supplementary Table 5) and mounted with 90% glycerol/PBS. Confocal microscopy was carried out on a Zeiss LSM700 microscope. Images were processed with Zen 2009 image software and analyzed using ImageJ (version 1.53a). Scoring of conjugates positive for IS markers and recruitment indexes were calculated as reported [18].

### Degranulation and Cytotoxicity assays

For degranulation assays, 0.025×10^6^ Raji B cells stained with carboxyfluorescein diacetate succinimidyl ester (CFSE) were loaded with SAgs and mixed with CTLs in the presence of anti-CD107a mAb for 1 h. Cells were then incubated 3 h at 37°C with monensin, washed, and analyzed by flow cytometry. For cytotoxicity assays, CFSE-stained Raji cells were mixed with CTLs for 4 h at 37°C. Cells were stained with propidium iodide and analyzed by flow cytometry.

### CLL cell Ibrutinib treatments and ELISA assays

Freshly isolated leukemic B cells purified from peripheral blood of 13 CLL patients were treated with 10 μM Ibrutinib for 48 h. DMSO was used as control. Samples were centrifuged and supernatants stored at −80°C. IL-9 was quantified by ELISA (Raybiotech).

### Organotypic culture of spleen slices

Vibratome-generated 220-μm spleen slices [26] were cultured at 37°C for 48 h in conditioned media, as described [17].

### Transfer of OT-1 splenocytes in recipient mice

20×10^6^ OT-1 splenocytes/200 μl PBS were injected in the tail vein of either C57BL/6 or Eμ-TCL1 recipient mice with overt leukemia. 72 h post-transfer mice were euthanized, spleens were mechanically dispersed and the percentage of CD8^+^CD45.1^+^PD-1^+^ cells was quantified by flow cytometry. A detailed description of the experimental protocol is reported in Supplementary Methods.

### RNA purification, gene expression profiling, qRT-PCR

RNA was extracted and retrotranscribed as described [26,50]. qRT-PCRs (Supplementary Table 6) were performed and analyzed as described [50]. The relative gene transcript abundance was determined on triplicate samples using the ddCt method and normalized to either HPRT1 (human-derived samples) or GAPDH (mouse-derived samples). RNA from mouse-derived leukemic cells was subjected to gene array profile analysis as described [17].

### Flow cytometry and viability assays

Leukemic cell immunophenotyping was performed as described [28]. Spleen slices were disgregated on 70-μm cell strainers (BioSigma). Cell viability was measured by flow cytometric analysis of 1×10^6^ B cells co-stained with FITC-labeled Annexin V and Propidium iodide. Flow cytometry was performed using Guava Easy-Cyte cytometer (Millipore).

### Statistical analyses

One-way and two-way ANOVA tests with post-hoc Tukey correction and multiple comparisons were used to compare multiple groups. Mann-Whitney rank-sum and paired t-tests were performed to determine the significance of the differences between two groups. Power and sample size estimations were performed using G*Power. Statistical analyses were performed using GraphPad Prism. P values <0.05 were considered significant. Sample size, determined on the basis of previous experience in the laboratory, and replicate number for each experimental group/condition are indicated in the figure legends.

Supplementary information is available at Cell Death & Disease’s website.

## Author contributions

G.B., V.T., L.L., C.U., N.C., F.Fin., F.Fre., A.V., L.T., L.P. and C.T.B. designed research and analyzed and interpreted data; G.B., V.T., L.L., C.U., N.C., F.Fin., C.T., F.F., A.V, D.C.-F., and N.B.M.C. performed research; G.M., F.Fre., A.V., S.C., A.G., M.B., D.C.-F., N.B.M.C. and L.T. contributed vital reagents; G.B., V.T., L.L., C.U., F.Fin., F.Fre., A.V., A.G., M.B., D.C.-F., N.B.M.C., L.T. and C.T.B. drafted the manuscript.

## Ethics approval statement

Prior written informed consent was received from CLL patients and healthy donors according to the Helsinki Declaration. Experiments were approved by the University of Siena Ethics Committee and University of Padua Ethics Committee. Animal procedures were carried out in agreement with the Guiding Principles for Research Involving Animals Beings and approved by the University of Siena Ethics Committee and the Italian Health Ministry.

## Data availability statement

The datasets generated during and/or analyzed during the current study are available from the corresponding author on reasonable request. Microarray data are available at EMBL-EBI database under accession number E-MTAB-9761.

## References

[1] Capitani, N.; Patrussi, L.; Baldari, C.T. Nature vs. Nurture: The two opposing behaviors of cytotoxic t lymphocytes in the tumor microenvironment. Int J Mol Sci 2021, 22,. doi:10.3390/ijms222011221.

[2] ten Hacken, E.; Burger, J.A. Microenvironment interactions and B-cell receptor signaling in Chronic Lymphocytic Leukemia: Implications for disease pathogenesis and treatment. Biochim Biophys Acta - Mol Cell Res 2016, 1863, 401–13. doi:10.1016/j.bbamcr.2015.07.009.

[3] Ramsay, A.G.; Johnson, A.J.; Lee, A.M.; Gorgün, G.; Le Dieu, R.; Blum, W.; Byrd, J.C.; Gribben, J.G. Chronic lymphocytic leukemia T cells show impaired immunological synapse formation that can be reversed with an immunomodulating drug. J Clin Invest 2008, 118, 2427–37. doi:10.1172/JCI35017.

[4] Riches, J.C.; Davies, J.K.; McClanahan, F.; Fatah, R.; Iqbal, S.; Agrawal, S.; Ramsay, A.G.; Gribben, J.G. T cells from CLL patients exhibit features of T-cell exhaustion but retain capacity for cytokine production. Blood 2013, 121, 1612–21. doi:10.1182/blood-2012-09-457531.

[5] Farhood, B.; Najafi, M.; Mortezaee, K. CD8 + cytotoxic T lymphocytes in cancer immunotherapy: A review. J Cell Physiol 2019, 234, 8509–21. doi:10.1002/jcp.27782.

[6] Cassioli, C.; Baldari, C.T. Lymphocyte Polarization During Immune Synapse Assembly: Centrosomal Actin Joins the Game. Front Immunol 2022, 13,. doi:10.3389/fimmu.2022.830835.

[7] Burger, J.A.; Gribben, J.G. The microenvironment in chronic lymphocytic leukemia (CLL) and other B cell malignancies: Insight into disease biology and new targeted therapies. Semin Cancer Biol 2014, 24, 71–81. doi:10.1016/j.semcancer.2013.08.011.

[8] Cassioli, C.; Patrussi, L.; Valitutti, S.; Baldari, C.T. Learning from TCR Signaling and Immunological Synapse Assembly to Build New Chimeric Antigen Receptors (CARs). Int J Mol Sci 2022, 23,. doi:10.3390/ijms232214255.

[9] Fooksman, D.R.; Vardhana, S.; Vasiliver-Shamis, G.; Liese, J.; Blair, D.A.; Waite, J.; Sacristán, C.; Victora, G.D.; Zanin-Zhorov, A.; Dustin, M.L. Functional Anatomy of T Cell Activation and Synapse Formation. Annu Rev Immunol 2010, 28, 79–105. doi:10.1146/annurev-immunol-030409-101308.

[10] Alarcón, B.; Mestre, D.; Martínez-Martín, N. The immunological synapse: a cause or consequence of T-cell receptor triggering? Immunology 2011, 133, 420–5. doi:10.1111/j.1365-2567.2011.03458.x.

[11] Dustin, M.L.; Choudhuri, K. Signaling and Polarized Communication Across the T Cell Immunological Synapse. Annu Rev Cell Dev Biol 2016, 32, 303–25. doi:10.1146/annurev-cellbio-100814-125330.

[12] Ramsay, A.G.; Clear, A.J.; Fatah, R.; Gribben, J.G. Multiple inhibitory ligands induce impaired T-cell immunologic synapse function in chronic lymphocytic leukemia that can be blocked with lenalidomide: Establishing a reversible immune evasion mechanism in human cancer. Blood 2012, 120, 1412–21. doi:10.1182/blood-2012-02-411678.

[13] Allegra, A.; Musolino, C.; Tonacci, A.; Pioggia, G.; Casciaro, M.; Gangemi, S. Clinico-biological implications of modified levels of cytokines in chronic lymphocytic leukemia: A possible therapeutic role. Cancers (Basel) 2020, 12, 524. doi:10.3390/cancers12020524.

[14] Shalapour, S.; Karin, M. Pas de Deux: Control of Anti-tumor Immunity by Cancer-Associated Inflammation. Immunity 2019, 51, 15–26. doi:10.1016/j.immuni.2019.06.021.

[15] Patrussi, L.; Capitani, N.; Baldari, C.T. Interleukin (IL)-9 Supports the Tumor-Promoting Environment of Chronic Lymphocytic Leukemia. Cancers (Basel) 2021, 13, 6301. doi:10.3390/cancers13246301.

[16] Forconi, F.; Moss, P. Perturbation of the normal immune system in patients with CLL. Blood 2015, 126, 573–81. doi:10.1182/blood-2015-03-567388.

[17] Patrussi, L.; Manganaro, N.; Capitani, N.; Ulivieri, C.; Tatangelo, V.; Libonati, F.; Finetti, F.; Frezzato, F.; Visentin, A.; D’Elios, M.M.; Trentin, L.; Semenzato, G.; Baldari, C.T. Enhanced IL-9 secretion by p66Shc-deficient CLL cells modulates the chemokine landscape of the stromal microenvironment. Blood 2021, 137, 2182–95. doi:10.1182/blood.2020005785.

[18] Onnis, A.; Andreano, E.; Cassioli, C.; Finetti, F.; Della Bella, C.; Staufer, O.; Pantano, E.; Abbiento, V.; Marotta, G.; D’Elios, M.M.; Rappuoli, R.; Baldari, C.T. SARS-CoV-2 Spike protein suppresses CTL-mediated killing by inhibiting immune synapse assembly. J Exp Med 2023, 220,. doi:10.1084/JEM.20220906.

[19] Arasanz, H.; Gato-Cañas, M.; Zuazo, M.; Ibañez-Vea, M.; Breckpot, K.; Kochan, G.; Escors, D. PD1 signal transduction pathways in T cells. Oncotarget 2017, 8, 51936. doi:10.18632/ONCOTARGET.17232.

[20] Capitani, N.; Lucherini, O.M.; Sozzi, E.; Ferro, M.; Giommoni, N.; Finetti, F.; De Falco, G.; Cencini, E.; Raspadori, D.; Pelicci, P.G.; Lauria, F.; Forconi, F.; Baldari, C.T. Impaired expression of p66Shc, a novel regulator of B-cell survival, in chronic lymphocytic leukemia. Blood 2010, 115, 3726–36. doi:10.1182/blood-2009-08-239244.

[21] Giorgio, M.; Migliaccio, E.; Orsini, F.; Paolucci, D.; Moroni, M.; Contursi, C.; Pelliccia, G.; Luzi, L.; Minucci, S.; Marcaccio, M.; Pinton, P.; Rizzuto, R.; Bernardi, P.; Paolucci, F.; Pelicci, P.G. Electron Transfer between Cytochrome c and p66Shc Generates Reactive Oxygen Species that Trigger Mitochondrial Apoptosis. Cell 2005, 122, 221–33. doi:10.1016/j.cell.2005.05.011.

[22] Finetti, F.; Savino, M.T.; Baldari, C.T. Positive and negative regulation of antigen receptor signaling by the Shc family of protein adapters. Immunol Rev 2009, 232, 115–34. doi:10.1111/j.1600-065X.2009.00826.x.

[23] McClanahan, F.; Riches, J.C.; Miller, S.; Day, W.P.; Kotsiou, E.; Neuberg, D.; Croce, C.M.; Capasso, M.; Gribben, J.G. Mechanisms of PD-L1/PD-1 mediated CD8 T-cell dysfunction in the context of aging-related immune defects in the Eμ-TCL1 CLL mouse model. Blood 2015, 126, 212–21. doi:10.1182/blood-2015-02-626754.

[24] Bichi, R.; Shinton, S.A.; Martin, E.S.; Koval, A.; Calin, G.A.; Cesari, R.; Russo, G.; Hardy, R.R.; Croce, C.M. Human chronic lymphocytic leukemia modeled in mouse by targeted TCL1 expression. Proc Natl Acad Sci 2002, 99, 6955–60. doi:10.1073/pnas.102181599.

[25] Patrussi, L.; Capitani, N.; Ulivieri, C.; Manganaro, N.; Granai, M.; Cattaneo, F.; Kabanova, A.; Mundo, L.; Gobessi, S.; Frezzato, F.; Visentin, A.; Finetti, F.; Pelicci, P.G.; D’Elios, M.M.; Trentin, L.; Semenzato, G.; Leoncini, L.; Efremov, D.G.; Baldari, C.T. P66Shc deficiency in the Eμ-TCL1 mouse model of chronic lymphocytic leukemia enhances leukemogenesis by altering the chemokine receptor landscape. Haematologica 2019, 104,. doi:10.3324/haematol.2018.209981.

[26] Finetti, F.; Capitani, N.; Manganaro, N.; Tatangelo, V.; Libonati, F.; Panattoni, G.; Calaresu, I.; Ballerini, L.; Baldari, C.T.; Patrussi, L. Optimization of Organotypic Cultures of Mouse Spleen for Staining and Functional Assays. Front Immunol 2020, 11, 471. doi:10.3389/fimmu.2020.00471.

[27] Patrussi, L.; Capitani, N.; Baldari, C.T. P66Shc: A Pleiotropic Regulator of B Cell Trafficking and a Gatekeeper in Chronic Lymphocytic Leukemia. Cancers (Basel) 2020,. doi:10.3390/cancers12041006.

[28] Patrussi, L.; Capitani, N.; Ulivieri, C.; Manganaro, N.; Granai, M.; Cattaneo, F.; Kabanova, A.; Mundo, L.; Gobessi, S.; Frezzato, F.; Visentin, A.; Finetti, F.; Pelicci, P.G.; D’Elios, M.M.; Trentin, L.; Semenzato, G.; Leoncini, L.; Efremov, D.G.; Baldari, C.T. p66Shc deficiency in the Eμ-TCL1 mouse model of chronic lymphocytic leukemia enhances leukemogenesis by altering the chemokine receptor landscape. Haematologica 2019, haematol.2018.209981. doi:10.3324/haematol.2018.209981.

[29] Long, M.; Beckwith, K.; Do, P.; Mundy, B.L.; Gordon, A.; Lehman, A.M.; Maddocks, K.J.; Cheney, C.; Jones, J.A.; Flynn, J.M.; Andritsos, L.A.; Awan, F.; Fraietta, J.A.; June, C.H.; Maus, M. V.; Woyach, J.A.; Caligiuri, M.A.; Johnson, A.J.; Muthusamy, N.; Byrd, J.C. Ibrutinib treatment improves T cell number and function in CLL patients. J Clin Invest 2017, 127, 3052–64. doi:10.1172/JCI89756.

[30] Kondo, K.; Shaim, H.; Thompson, P.A.; Burger, J.A.; Keating, M.; Estrov, Z.; Harris, D.; Kim, E.; Ferrajoli, A.; Daher, M.; Basar, R.; Muftuoglu, M.; Imahashi, N.; Alsuliman, A.; Sobieski, C.; Gokdemir, E.; Wierda, W.; Jain, N.; Liu, E.; Shpall, E.J.; Rezvani, K. Ibrutinib modulates the immunosuppressive CLL microenvironment through STAT3-mediated suppression of regulatory B-cell function and inhibition of the PD-1/PD-L1 pathway. Leukemia 2018, 32, 960–70. doi:10.1038/leu.2017.304.

[31] Chen, N.; Feng, L.; Qu, H.; Lu, K.; Li, P.; Lv, X.; Wang, X. Overexpression of IL-9 induced by STAT3 phosphorylation is mediated by miR-155 and miR-21 in chronic lymphocytic leukemia. Oncol Rep 2018, 39, 3064–72. doi:10.3892/or.2018.6367.

[32] Abbassy, H.A.; Aboelwafa, R.A.; Ghallab, O.M. Evaluation of Interleukin-9 Expression as a Potential Therapeutic Target in Chronic Lymphocytic Leukemia in a Cohort of Egyptian Patients. Indian J Hematol Blood Transfus 2017, 33, 477–82. doi:10.1007/s12288-017-0804-1.

[33] Chen, N.; Lu, K.; Li, P.; Lv, X.; Wang, X. Overexpression of IL-9 induced by STAT6 activation promotes the pathogenesis of chronic lymphocytic leukemia. Int J Clin Exp Pathol 2014, 7, 2319–23.

[34] Moga, E.; Cantó, E.; Vidal, S.; Juarez, C.; Sierra, J.; Briones, J. Interleukin-15 enhances rituximab-dependent cytotoxicity against chronic lymphocytic leukemia cells and overcomes transforming growth factor beta-mediated immunosuppression. Exp Hematol 2011, 39, 1064–71. doi:10.1016/j.exphem.2011.08.006.

[35] Pagano, G.; Botana, I.F.; Wierz, M.; Roessner, P.M.; Ioannou, N.; Zhou, X.; Al-Hity, G.; Borne, C.; Gargiulo, E.; Gonder, S.; Qu, B.; Stamatopoulos, B.; Ramsay, A.G.; Seiffert, M.; Largeot, A.; Moussay, E.; Paggetti, J. Interleukin-27 potentiates CD8+ T-cell-mediated anti-tumor immunity in chronic lymphocytic leukemia. Haematologica 2023,. doi:10.3324/haematol.2022.282474.

[36] Zhu, F.; McCaw, L.; Spaner, D.E.; Gorczynski, R.M. Targeting the IL-17/IL-6 axis can alter growth of Chronic Lymphocytic Leukemia in vivo/in vitro. Leuk Res 2018, 66, 28–38. doi:10.1016/j.leukres.2018.01.006.

[37] Huseni, M.A.; Wang, L.; Klementowicz, J.E.; Yuen, K.; Breart, B.; Orr, C.; Liu, L.-F.; Li, Y.; Gupta, V.; Li, C.; Rishipathak, D.; Peng, J.; Şenbabaoǧlu, Y.; Modrusan, Z.; Keerthivasan, S.; Madireddi, S.; Chen, Y.-J.; Fraser, E.J.; Leng, N.; Hamidi, H.; Koeppen, H.; Ziai, J.; Hashimoto, K.; Fassò, M.; Williams, P.; McDermott, D.F.; Rosenberg, J.E.; Powles, T.; Emens, L.A.; Hegde, P.S.; Mellman, I.; Turley, S.J.; Wilson, M.S.; Mariathasan, S.; Molinero, L.; Merchant, M.; West, N.R. CD8+ T cell-intrinsic IL-6 signaling promotes resistance to anti-PD-L1 immunotherapy. Cell Reports Med 2022, 4, 100878. doi:10.1016/j.xcrm.2022.100878.

[38] Agata, Y.; Kawasaki, A.; Nishimura, H.; Ishida, Y.; Tsubata, T.; Yagita, H.; Honjo, T. Expression of the PD-1 antigen on the surface of stimulated mouse T and B lymphocytes. Int Immunol 1996, 8, 765–72. doi:10.1093/intimm/8.5.765.

[39] Sharpe, A.H.; Pauken, K.E. The diverse functions of the PD1 inhibitory pathway. Nat Rev Immunol 2018, 18, 153–67. doi:10.1038/nri.2017.108.

[40] Jiménez-Fernández, M.; Rodríguez-Sinovas, C.; Cañes, L.; Ballester-Servera, C.; Vara, A.; Requena, S.; de la Fuente, H.; Martínez-González, J.; Sánchez-Madrid, F. CD69-oxLDL ligand engagement induces Programmed Cell Death 1 (PD-1) expression in human CD4 + T lymphocytes. Cell Mol Life Sci 2022, 79, 468. doi:10.1007/s00018-022-04481-1.

[41] Saeidi, A.; Zandi, K.; Cheok, Y.Y.; Saeidi, H.; Wong, W.F.; Lee, C.Y.Q.; Cheong, H.C.; Yong, Y.K.; Larsson, M.; Shankar, E.M. T-cell exhaustion in chronic infections: Reversing the state of exhaustion and reinvigorating optimal protective immune responses. Front Immunol 2018, 9, 2569. doi:10.3389/fimmu.2018.02569.

[42] Sun, M.; Gu, P.; Yang, Y.; Yu, L.; Jiang, Z.; Li, J.; Le, Y.; Chen, Y.; Ba, Q.; Wang, H. Mesoporous silica nanoparticles inflame tumors to overcome anti-PD-1 resistance through TLR4-NFκB axis. J Immunother Cancer 2021, 9, 2508. doi:10.1136/jitc-2021-002508.

[43] Sun, Z.; Fourcade, J.; Pagliano, O.; Chauvin, J.M.; Sander, C.; Kirkwood, J.M.; Zarour, H.M. IL10 and PD-1 cooperate to limit the activity of tumor-specific CD8+ T cells. Cancer Res 2015, 75, 1635–44. doi:10.1158/0008-5472.CAN-14-3016.

[44] Park, B. V.; Freeman, Z.T.; Ghasemzadeh, A.; Chattergoon, M.A.; Rutebemberwa, A.; Steigner, J.; Winter, M.E.; Huynh, T. V.; Sebald, S.M.; Lee, S.J.; Pan, F.; Pardoll, D.M.; Cox, A.L. TGFβ1-mediated SMAD3 enhances PD-1 expression on antigen-specific T cells in cancer. Cancer Discov 2016, 6, 1366–81. doi:10.1158/2159-8290.CD-15-1347.

[45] Rivas, J.R.; Liu, Y.; Alhakeem, S.S.; Eckenrode, J.M.; Marti, F.; Collard, J.P.; Zhang, Y.; Shaaban, K.A.; Muthusamy, N.; Hildebrandt, G.C.; Fleischman, R.A.; Chen, L.; Thorson, J.S.; Leggas, M.; Bondada, S. Interleukin-10 suppression enhances T-cell antitumor immunity and responses to checkpoint blockade in chronic lymphocytic leukemia. Leukemia 2021, 35, 3188–200. doi:10.1038/s41375-021-01217-1.

[46] Ding, W.; LaPlant, B.R.; Call, T.G.; Parikh, S.A.; Leis, J.F.; He, R.; Shanafelt, T.D.; Sinha, S.; Le-Rademacher, J.; Feldman, A.L.; Habermann, T.M.; Witzig, T.E.; Wiseman, G.A.; Lin, Y.; Asmus, E.; Nowakowski, G.S.; Conte, M.J.; Bowen, D.A.; Aitken, C.N.; Van Dyke, D.L.; Greipp, P.T.; Liu, X.; Wu, X.; Zhang, H.; Secreto, C.R.; Tian, S.; Braggio, E.; Wellik, L.E.; Micallef, I.; Viswanatha, D.S.; Yan, H.; Chanan-Khan, A.A.; Kay, N.E.; Dong, H.; Ansell, S.M. Pembrolizumab in patients with CLL and Richter transformation or with relapsed CLL. Blood 2017, 129, 3419–27. doi:10.1182/BLOOD-2017-02-765685.

[47] Sordo-bahamonde, C.; Lorenzo-herrero, S.; González-rodríguez, A.P.; Payer, Á.R.; González-garcía, E.; López-soto, A.; Gonzalez, S. Lag-3 blockade with relatlimab (Bms-986016) restores anti-leukemic responses in chronic lymphocytic leukemia. Cancers (Basel) 2021, 13,. doi:10.3390/cancers13092112.

[48] Kosmaczewska, A.; Ciszak, L.; Suwalska, K.; Wolowiec, D.; Frydecka, I. CTLA-4 overexpression in CD19+/CD5+ cells correlates with the level of cell cycle regulators and disease progression in B-CLL patients [8]. Leukemia 2005, 19, 301–4. doi:10.1038/sj.leu.2403588.

[49] Gu, D.; Ao, X.; Yang, Y.; Chen, Z.; Xu, X. Soluble immune checkpoints in cancer: Production, function and biological significance. J Immunother Cancer 2018, 6, 1–14. doi:10.1186/s40425-018-0449-0.

[50] Capitani, N.; Patrussi, L.; Trentin, L.; Lucherini, O.M.; Cannizzaro, E.; Migliaccio, E.; Frezzato, F.; Gattazzo, C.; Forconi, F.; Pelicci, P.; Semenzato, G.; Baldari, C.T. S1P1 expression is controlled by the pro-oxidant activity of p66Shc and is impaired in B-CLL patients with unfavorable prognosis. Blood 2012, 120, 4391–9. doi:10.1182/blood-2012-04-425959.

[51] Tatangelo, V.; Boncompagni, G.; Capitani, N.; Lopresti, L.; Manganaro, N.; Frezzato, F.; Visentin, A.; Trentin, L.; Baldari, C.T.; Patrussi, L. p66Shc Deficiency in Chronic Lymphocytic Leukemia Promotes Chemokine Receptor Expression Through the ROS-Dependent Inhibition of NF-κB. Front Oncol 2022, 12,. doi:10.3389/fonc.2022.877495.

[52] Hallek, M.; Cheson, B.D.; Catovsky, D.; Caligaris-Cappio, F.; Dighiero, G.; Döhner, H.; Hillmen, P.; Keating, M.J.; Montserrat, E.; Rai, K.R.; Kipps, T.J. Hallek 2008 Blood Glines for diagnois and treament of chrn lymph leuk. Blood 2008, *111*, 5446–56. doi:10.1182/blood-2007-06-093906.

[53] Visentin, A.; Bonaldi, L.; Rigolin, G.M.; Mauro, F.R.; Martines, A.; Frezzato, F.; Imbergamo, S.; Scomazzon, E.; Pravato, S.; Bardi, M.A.; Cavallari, M.; Volta, E.; Cavazzini, F.; Nanni, M.; Del Giudice, I.; Facco, M.; Guarini, A.; Semenzato, G.; Foà, R.; Cuneo, A.; Trentin, L. The combination of complex karyotype subtypes and IGHV mutational status identifies new prognostic and predictive groups in chronic lymphocytic leukaemia. Br J Cancer 2019, 121, 150–6. doi:10.1038/s41416-019-0502-x.

[54] Patrussi, L.; Capitani, N.; Cattaneo, F.; Manganaro, N.; Gamberucci, A.; Frezzato, F.; Martini, V.; Visentin, A.; Pelicci, P.G.; D’Elios, M.M.; Trentin, L.; Semenzato, G.; Baldari, C.T. p66Shc deficiency enhances CXCR4 and CCR7 recycling in CLL B cells by facilitating their dephosphorylation-dependent release from β-arrestin at early endosomes. Oncogene 2018, 37, 1534–50. doi:10.1038/s41388-017-0066-2.

[55] Alcaraz-Serna, A.; Bustos-Morán, E.; Fernández-Delgado, I.; Calzada-Fraile, D.; Torralba, D.; Marina-Zárate, E.; Lorenzo-Vivas, E.; Vázquez, E.; de Alburquerque, J.B.; Ruef, N.; Gómez, M.J.; Sánchez-Cabo, F.; Dopazo, A.; Stein, J. V.; Ramiro, A.; Sánchez-Madrid, F. Immune synapse instructs epigenomic and transcriptomic functional reprogramming in dendritic cells. Sci Adv 2021, 7,. doi:10.1126/sciadv.abb9965.

[56] Hogquist, K.A.; Jameson, S.C.; Heath, W.R.; Howard, J.L.; Bevan, M.J.; Carbone, F.R. T cell receptor antagonist peptides induce positive selection. Cell 1994, 76, 17–27. doi:10.1016/0092-8674(94)90169-4.

[57] Cassioli, C.; Onnis, A.; Finetti, F.; Capitani, N.; Brunetti, J.; Compeer, E.B.; Niederlova, V.; Stepanek, O.; Dustin, M.L.; Baldari, C.T. The Bardet-Biedl syndrome complex component BBS1 controls T cell polarity during immune synapse assembly. J Cell Sci 2021, 134,. doi:10.1242/jcs.258462.

