## Supplementary Methods, Figures and Tables for "Leukemic cell-secreted interleukin-9 suppresses cytotoxic T cell-mediated killing in chronic lymphocytic leukemia"

#### Purification, activation and conditioning of $CD8^+$ cells

$CD8^+$ T cells isolated from spleens of C57BL/6J and Eμ-TCL1 by immunomagnetic sorting using Dynabeads™ Untouched Mouse CD8 Cells Kit were seeded on 6-well plates ($1.5x10^5$ cells/well) and incubated for 48 h with 2 ml supernatant conditioned by either leukemic or wild-type B cells [1]. 0.5 ng/ml recombinant murine IL-9 and 0.1 ng/ml of either control or anti-IL-9 mAb (Supplementary Table 4) were added to culture media.

$CD8^+$ cells were isolated from peripheral blood of healthy donors and CLL patients by negative selection using the RosetteSep Human $CD8^+$ T Cell Enrichment Cocktail, following the manufacturer's instructions. On the same day (day 0), cells were stimulated in RPMI-HEPES medium ($1\times10^6$/ml) (#R7388; Merck) supplemented with 10% BCS (#SH30072.03; GE Healthcare HyClone), 1% MEM nonessential amino acids (MEM NEAA; #11140050), and 50 U/ml recombinant human IL-2 with Dynabeads™ Human T-activator CD3/CD28. Media conditioned by either leukemic or wild-type B cells were immediately added to the activation medium, to a final 1:1 ratio (conditioned *vs* activation medium). 20 ng/ml recombinant human IL-9 or 0.5 ng/ml recombinant human IL-10, and 0.1 ng/ml isotype control, 1 ng/ml anti-IL-9 mAb or anti-IL10 mAb (Supplementary Table 4), were added to culture media. 48 h after activation (day 2), beads were removed and CTLs were collected. For cytotoxicity and degranulation assays, CTLs were expanded in RPMI-HEPES supplemented with 10% BCS, 1% MEM NEAA, and 50 U/ml recombinant human IL-2 for additional 3 days, then further expanded for 2 days and collected at day 7 [2].

#### In vivo Ibrutinib treatment of CLL patients

PB was collected from 4 CLL patients treated first line with chemoimmunotherapy. Patients from #1 to #3 received FCR (fludarabine 25 mg/ml plus cyclophosphamide 250 mg/ml administered on day 1-3 of cycles 1-6 and rituximab 375 mg/m$^2$ on day 1 of cycle 1 and 500 mg/ml on day 1 of cycles 2-6). Patient #4 received BR (bendamustine 90 mg/m$^2$ administered on day 1-2 of cycles 1-6 and rituximab 375 mg/m$^2$ on day 1 of cycle 1 and 500 mg/ml on day 1 of cycles 2-6). These treatments were started at disease progression according to iwCLL criteria [3]. At disease relapse all patients were managed with ibrutinib 420 mg once a day [4]. From each patient, PB samples were collected the starting day of ibrutinib treatment and during follow-up [3,4].

**Immune synapse formation, immunofluorescence acquisition and analysis**

Raji B cells (0.4×10$^6$ cells/100 μl) were loaded with 10 μg/ml Staphylococcal SAg A (SEA; Toxin Technologies, #AT101), B (SEB; Toxin Technologies, #BT202) and E (SEE; Toxin Technologies, #ET404) for 2 h to broadly cover the TCR Vβ repertoire, and labelled with 20 μM Cell Tracker Blue for 15 min [2]. Conjugates of CTLs with Raji B cells formed in the absence of SAgs were used as negative controls. Raji B cells were mixed with CTLs (1:1.5) and incubated for 15 min at 37°C. When required, 7.5 μg/ml anti-PD-1 neutralizing mAbs was added to the medium during conjugate formation. Samples were seeded onto poly-L-lysine (Merck, #P1274)-coated slides (ThermoFisher Scientific, #X2XER208B), fixed for 10 min in methanol at -20°C (for CD3ζ and pTyr staining) or for 15 min with 4% paraformaldehyde/PBS at room temperature (for PCNT and phalloidin staining) and permeabilized with 0.1% Triton, 1% BSA PBS. Cells were stained with primary antibodies overnight at 4°C, washed with PBS, incubated for 45 min at room temperature with Alexa Fluor 488- and 555-labeled secondary antibodies and mounted with 90% glycerol/PBS. Confocal microscopy was carried out on a Zeiss LSM700 (Carl Zeiss, Jena, Germany) microscope using a 63x/1.40 objective. Images were acquired with pinholes opened to obtain 0.8 μm-tick sections. Images were processed with Zen 2009 image software (Carl Zeiss, Jena, Germany). Immunofluorescence analyses were performed using ImageJ (RRID:SCR_003070). Scoring of conjugates for accumulation of CD3ζ, p-Tyr or F-actin at the IS, or for centrosome (PCNT) juxtaposition to the IS membrane, was performed as reported [2,5]. Recruitment indexes and quantification of the relative distances (μm) of the centrosome (PCNT staining) from the center of the contact site with the APC were calculated using ImageJ [2,5]. Antibodies used for immunofluorescence microscopy are listed in Supplementary Table 4.

**Degranulation and cytotoxicity assays**

For degranulation assays [2], Raji B cells were incubated with 1.5 μM carboxyfluorescein diacetate succinimidyl ester (CFSE; #C34554; Thermo Fisher Scientific) dissolved in PBS for 8 min at room temperature. CFSE-stained Raji B cells ($0.025\times10^6$) were pulsed with 1 μg/ml SAgs for 1 h in serum-free AIMV medium (#12055-091; Gibco), then mixed with CTLs, at the ratios of 1:2.5, 1:5 and 1:10 (APC:CTL ratio) in 50 μl AIMV medium containing APC-labeled anti-human CD107a (LAMP1; BioLegend) mAb for 1 h. Monensin (BioLegend) was then added and cells were further incubated for further 3 h at 37 °C. Unpulsed CFSE-stained Raji B cells were used as negative control. Then cells were washed, resuspended in cold PBS and acquired using a GUAVA flow cytometer (Merck Millipore).

For cytotoxicity assays [2], Raji B cells ($0.025\times10^6$) were stained with 1.5 μM CFSE; (#C34554; Thermo Fisher Scientific) 8 min at room temperature in PBS and then pulsed with 2 μg/ml SAgs for 1 h in serum-free AIMV medium. Unpulsed CFSE-stained Raji B cells were used as negative control. CTLs collected at day 7 were added to Raji B cells at different ratios (1:2.5, 1:5 and 1:10 APC:CTL ratio) in 50 μl AIMV medium and incubated at

37°C for 4 h to evaluate target cell killing. Raji B cells w/o SAgs were used to set up control samples. Cells were then diluted to 200 μl with cold PBS and acquired using a GUAVA flow cytometer. Propidium Iodide (PI, Sigma, #537059) was added before each acquisition to the final concentration of 20 μg/ml. Cytotoxicity (% target cell lysis) was calculated as follows: $CFSE^+PI^+$ cells − $CFSE^+PI^+$ cells in control sample) × 100.

**Adoptive transfer of splenocytes from OT-1 mice**

OT-1 TCR transgenic splenocytes were harvested from the spleen of 5 naive CD45.1 OT-1 mice by mechanical dissociation using a 40 μm cell strainer (Falcon™, Thermo Fisher Scientific) into a petri dish rinsed with complete medium. Cells were pooled and resuspended in PBS at $10^8$ cells/ml before the adoptive transfer. 20 × $10^6$ OT-1 splenocytes were injected in the tail vein (200 μl/mice) of CD45.2 Eμ-TCL1 recipient mice with overt leukemia (~40% $CD5^+CD19^+$ leukemic cells in PB) (n=4) or age-matched C57BL/6 mice (n=4). 72 h after adoptive transfer mice were euthanized, spleens were homogenized and cells were stained with anti-CD8, anti-CD45.1 and anti-PD-1 antibodies. The percentage of $CD8^+$ $PD\text{-}1^+$ cells was quantified by flow cytometry on $CD45.1^+$ positive cells.

### References

1. Patrussi L, Manganaro N, Capitani N, Ulivieri C, Tatangelo V, Libonati F, et al. Enhanced IL-9 secretion by p66Shc-deficient CLL cells modulates the chemokine landscape of the stromal microenvironment. Blood. 2021; 137(16): 2182–95.

2. Onnis A, Andreano E, Cassioli C, Finetti F, Della Bella C, Staufer O, et al. SARS-CoV-2 Spike protein suppresses CTL-mediated killing by inhibiting immune synapse assembly. J Exp Med [Internet]. 2023 [cited 2022 Dec 13]; 220(2). Available from: /pmc/articles/PMC9671159/

3. Eichhorst B, Fink A-M, Bahlo J, Busch R, Kovacs G, Maurer C, et al. First-line chemoimmunotherapy with bendamustine and rituximab versus fludarabine, cyclophosphamide, and rituximab in patients with advanced chronic lymphocytic leukaemia (CLL10): an international, open-label, randomised, phase 3, non-inferiority trial. Lancet Oncol [Internet]. 2016; 17(7): 928–42. Available from: https://linkinghub.elsevier.com/retrieve/pii/S1470204516300511

4. Brown JR, Hillmen P, O'Brien S, Barrientos JC, Reddy NM, Coutre SE, et al. Extended follow-up and impact of high-risk prognostic factors from the phase 3 RESONATE study in patients with previously treated CLL/SLL. Leukemia [Internet]. 2018; 32(1): 83–91. Available from: http://www.nature.com/doifinder/10.1038/leu.2017.175

5. Cassioli C, Onnis A, Finetti F, Capitani N, Brunetti J, Compeer EB, et al. The Bardet-Biedl syndrome complex component BBS1 controls T cell polarity during immune synapse assembly. J Cell Sci [Internet]. 2021 [cited 2021 Sep 10]; 134(16). Available from: https://pubmed.ncbi.nlm.nih.gov/34251457/

### Supplementary Figures

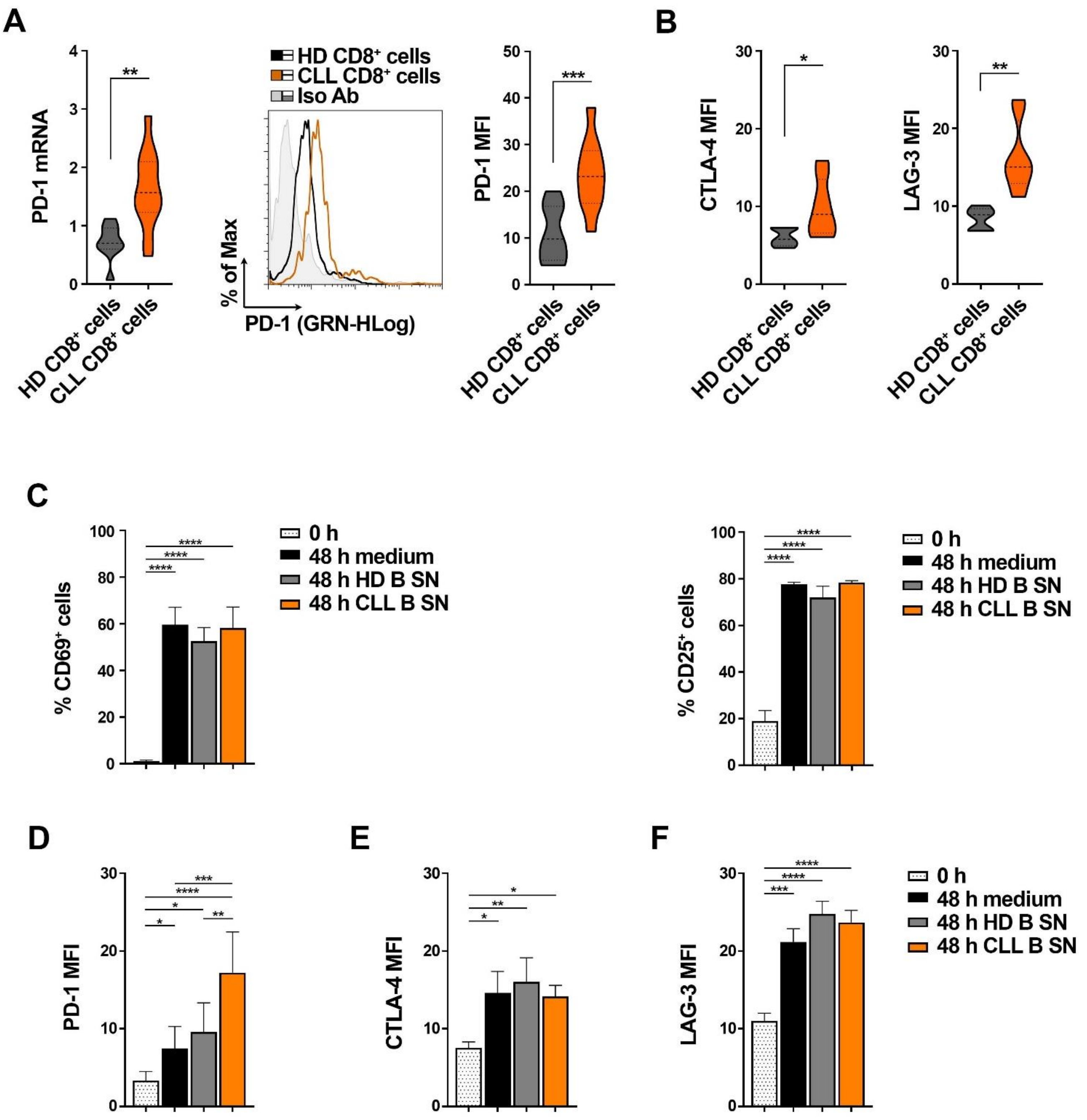


Figure S1

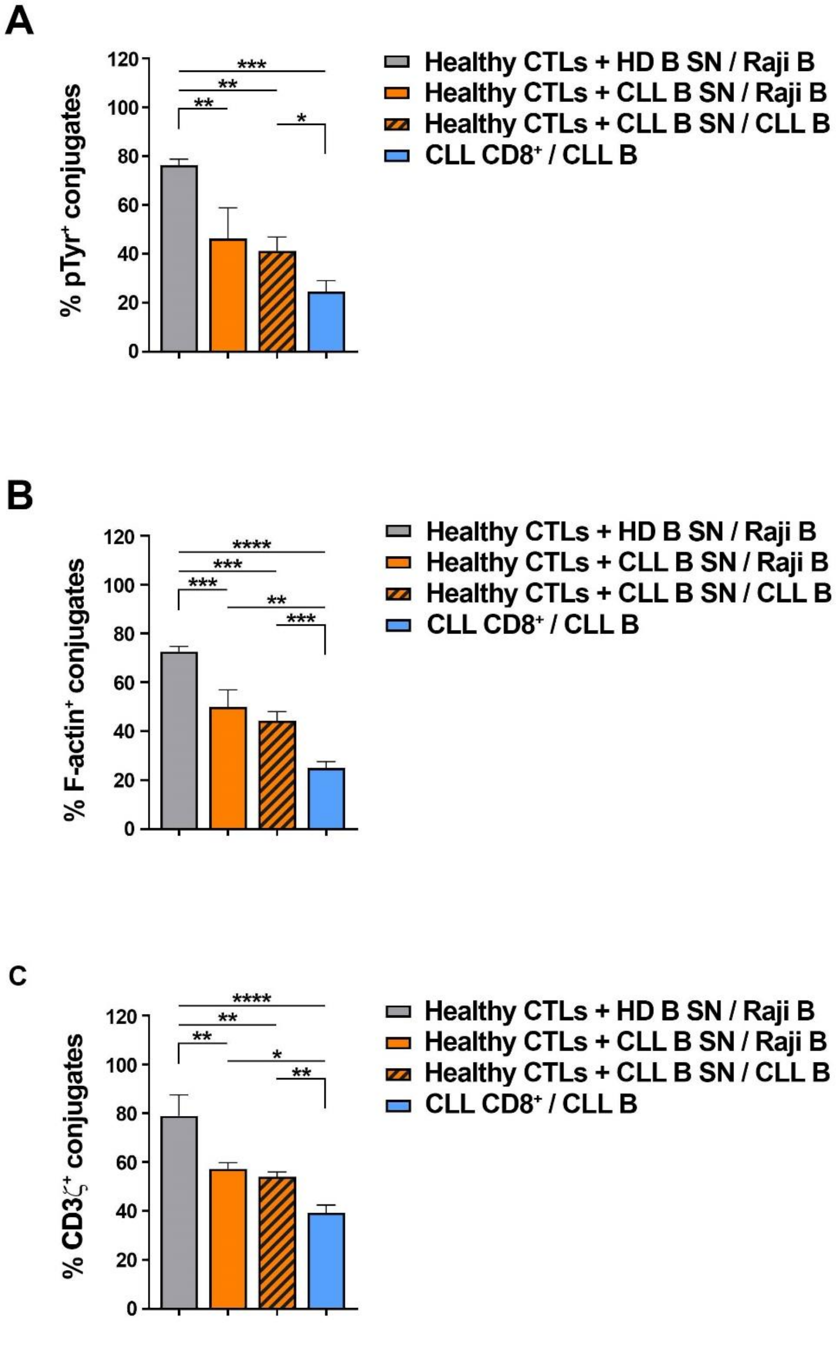
A
% pTyr+ conjugates
0
20
40
60
80
100
120
***
**
**
*
Healthy CTLs + HD B SN / Raji B
Healthy CTLs + CLL B SN / Raji B
Healthy CTLs + CLL B SN / CLL B
CLL CD8+ / CLL B
B
% F-actin+ conjugates
0
20
40
60
80
100
120
****
***
***
**
***
Healthy CTLs + HD B SN / Raji B
Healthy CTLs + CLL B SN / Raji B
Healthy CTLs + CLL B SN / CLL B
CLL CD8+ / CLL B
C
% CD3ζ+ conjugates
0
20
40
60
80
100
120
****
**
**
*
**
Healthy CTLs + HD B SN / Raji B
Healthy CTLs + CLL B SN / Raji B
Healthy CTLs + CLL B SN / CLL B
CLL CD8+ / CLL B

**Figure S2**

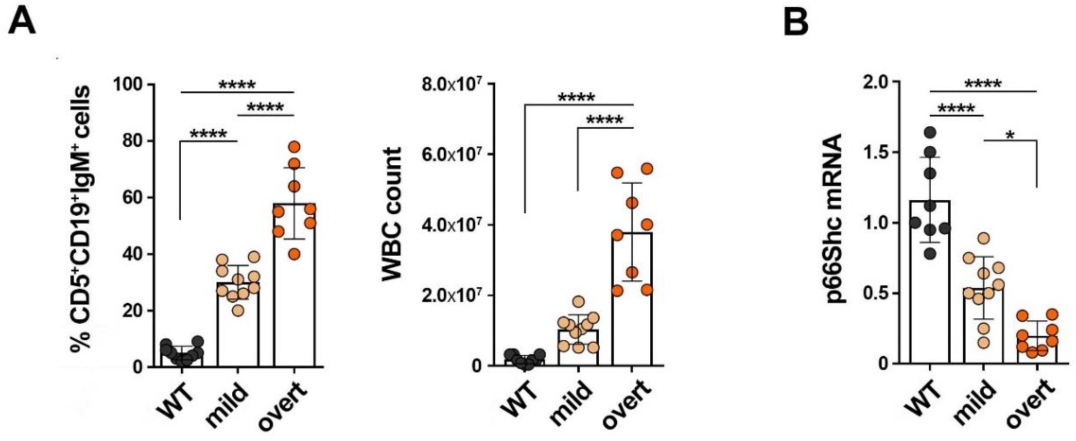

A
B
% CD5+CD19+IgM+ cells
100
80
60
40
20
0
****
WT
mild
overt
WBC count
8.0x10^7
6.0x10^7
4.0x10^7
2.0x10^7
p66Shc mRNA
2.0
1.5
1.0
0.5
*


Figure S3

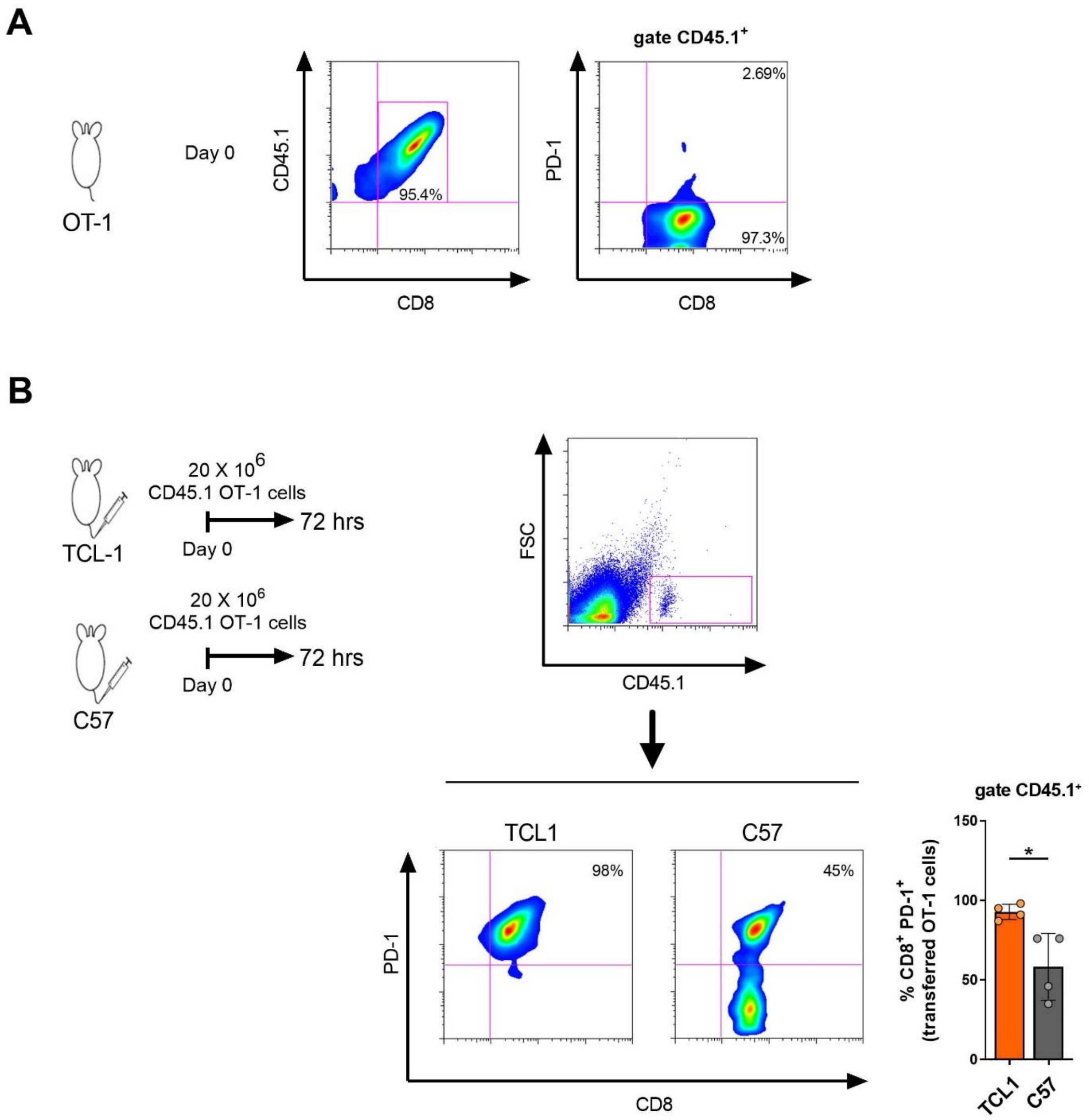
A
OT-1
Day 0
CD45.1
95.4%
CD8
gate CD45.1+
PD-1
2.69%
97.3%
CD8
B
TCL-1
20 X 10^6
CD45.1 OT-1 cells
72 hrs
Day 0
C57
20 X 10^6
CD45.1 OT-1 cells
72 hrs
Day 0
FSC
CD45.1
TCL1
C57
98%
45%
PD-1
CD8
gate CD45.1+
% CD8+ PD-1+ (transferred OT-1 cells)
150
100
50
0
*
TCL1
C57

Figure S4

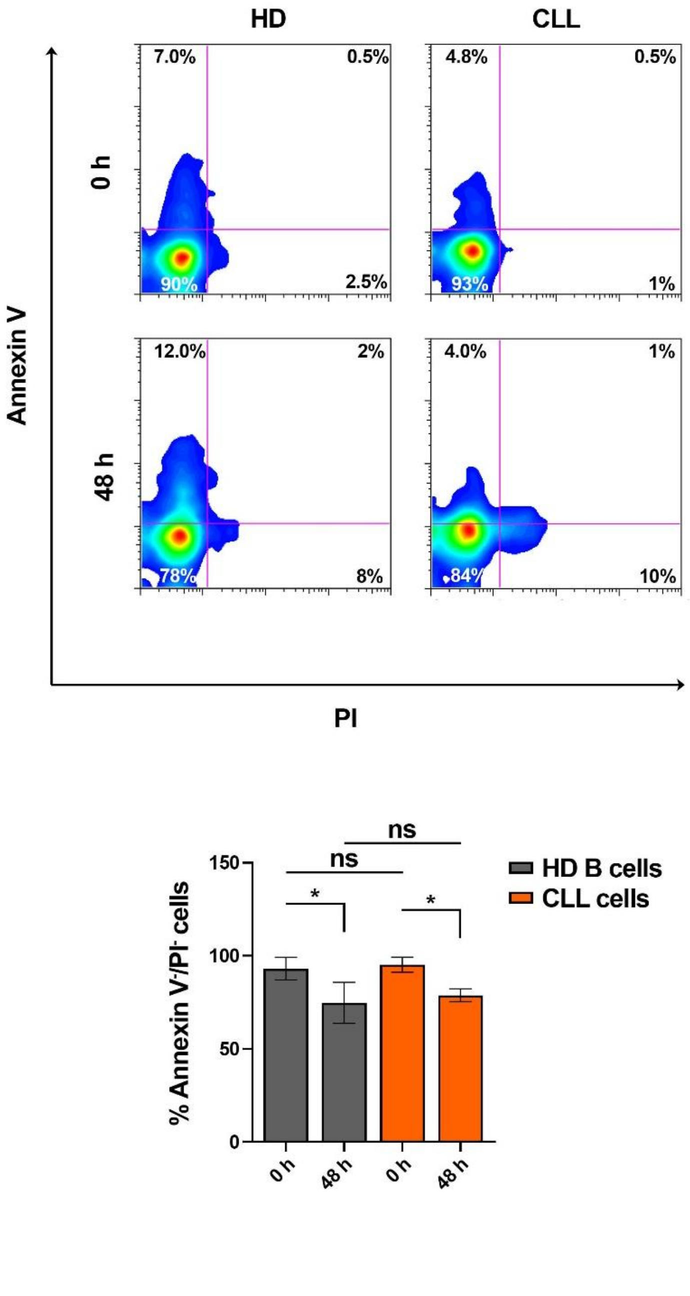
HD
CLL
0 h
48 h
7.0%
0.5%
90%
2.5%
4.8%
0.5%
93%
1%
12.0%
2%
78%
8%
4.0%
1%
84%
10%
Annexin V
PI
ns
ns
*
*
150
100
50
0
% Annexin V-/PI- cells
HD B cells
CLL cells
0 h
48 h
0 h
48 h

**Figure S5**

### Legends to Supplementary Figures

**Figure S1. A.** qRT-PCR analysis of mRNA (*left*) and flow cytometric analysis of surface expression (*right*) of PD-1 in $CD8^+$ cells purified from PB of healthy donors ($n \geq 8$) or CLL patients ($n \geq 10$). The flow cytometric histogram shows PD-1 staining in $CD8^+$ cells from a representative healthy donor and a representative CLL patient. The relative gene transcript abundance was determined on triplicate samples using the ddCt method and normalized to HPRT1 (Mann-Whitney Rank Sum test). **B.** Flow cytometric analysis of surface expression of CTLA-4 (*left*) and LAG-3 (*right*) in $CD8^+$ cells purified from PB of healthy donors (n = 5) or CLL patients (n = 5). **C**. Flow cytometric analysis of surface expression of CD69 (left) or CD25 (right) in $CD8^+$ cells purified from PB of healthy donors (n = 3), either immediately after purification (0 h) or activated for 48 h in the presence of media conditioned by healthy B cells / B cells purified from CLL patients. (n = 3). **D-F**. Flow cytometric analysis of surface expression of PD-1 (*left*), CTLA-4 (*middle*) and LAG-3 (*right*) in $CD8^+$ cells purified from PB of healthy donors (n = 3), either immediately after purification (0 h) or activated for 48 h in the presence of complete culture medium (48 h medium) or media conditioned by healthy B cells (n = 7) / B cells purified from CLL patients (n = 8). (**A, B**: Mann-Whitney Rank Sum test; **C-F**: Ordinary one-way ANOVA test) ****, $p \leq 0.0001$; ***, $p \leq 0.001$; **, $p \leq 0.01$; *, $p \leq 0.05$.

**Figure S2**. Immunofluorescence analysis of pTyr (**A**), F-actin (**B**), and CD3ζ (**C**) in either CTLs activated for 48 h in the presence of media conditioned by healthy B cells / B cells purified from CLL patients, or T cells purified from CLL patients (CLL T), mixed with either Raji cells (Raji B) or B cells purified from CLL patients (CLL B). B cells were pulsed with a combination of SEA, SEB, and SEE (SAgs), and incubated for 15 min at 37°C prior to mixing with T cells. Data are expressed as % of 15-min SAg-specific conjugates harboring staining at the IS (≥50 cells/sample, n = 3, Ordinary one-way ANOVA test). ****, $p \leq 0.0001$; ***, $p \leq 0.001$; **, $p \leq 0.01$; *, $p \leq 0.05$.

**Figure S3.** **A**. Flow cytometric analysis of the % of $CD19^+CD5^+IgM^+$ cells (left) and quantification of the number of white blood cells (WBC) (right) in PB of wild type (WT, n = 9) and Eμ-TCL1 mice with mild (n = 10) or overt (n = 8) leukemia (Ordinary one-way ANOVA). **B**. qRT-PCR analysis of p66Shc mRNA in B cells purified from spleens of wild type (WT, n = 8) and Eμ-TCL1 mice with mild (n = 10) or overt (n = 8) leukemia (Ordinary one-way ANOVA). The relative gene transcript abundance was determined on triplicate samples using the ddCt method and normalized to GAPDH. ****, $p \leq 0.0001$; *, $p \leq 0.05$.

**Figure S4. A.** Flow cytometric analysis of the percentage of $CD8^+$ PD-$1^+$ CD45.$1^+$ cells in splenocytes of OT-1 mice. Splenocytes were pooled from 5 spleens and stained immediately before adoptive transfer in recipient mice. **B.** Flow cytometric analysis of the percentage of $CD8^+$ PD-$1^+$ CD45.$1^+$ cells in splenocytes obtained from wild-type (C57, n = 4) or Eμ-TCL1 (TCL1, n = 4) mice injected with 20 x $10^6$ OT-1 splenocytes in 200 μl PBS. Recipient mice were euthanized 72 h after adoptive transfer. Results are shown as the percentage of $CD8^+$ PD-$1^+$ cells on CD45.$1^+$ cells. Gating strategy and representative flow cytometric plots are shown. Paired t test; *, $p \leq 0.05$.

**Figure S5.** Flow cytometric analysis of the percentage of viable cells in B cells purified from peripheral blood of healthy donors (healthy B cells, n donors = 3) or CLL patients (CLL cells, n CLL patients = 3), either immediately after purification (0 h) or after culture for 48 h (48 h) in complete culture media and staining with Annexin V/PI. Representative flow cytometric plots are shown. Mean±SD. ns: not significant. (Ordinary one-way ANOVA). $p \leq 0.05$, *.

### Supplementary Tables

**Supplementary Table 1. p66Shc deficiency in Eμ-TCL1 mice correlates with enhanced secretion of CCL22, CCL24, IL-9 and IL-10 by leukemic cells.** List and concentrations (ng/ml) of the soluble factors released in the culture supernatants of either B cells purified from wild-type mice (n = 7) or leukemic cells purified from Eμ-TCL1 (TCL1, n = 15) and Eμ-TCL1/p66Shc$^{-/-}$ (TCL1/p66$^{-/-}$, n = 16) mice. The amounts of CCL22, CCL24, IL-9 and IL-10 (highlighted in bold) were significantly enhanced in Eμ-TCL1 and Eμ-TCL1/p66Shc$^{-/-}$ *vs* wild-type supernatants, and in Eμ-TCL1/p66$^{-/-}$ *vs* Eμ-TCL1. (Student's t test). ***, $p \leq 0.001$; **, $p \leq 0.01$; *, $p \leq 0.05$.

| Protein | C57BL/6 SN ng/ml | TCL1 SN ng/ml | TCL1/p66$^{-/-}$ SN ng/ml | *p*-value TCL1 SN *vs* C57BL/6 SN | *p*-value TCL1/p66$^{-/-}$ SN *vs* C57BL/6 SN | *p*-value TCL1/p66$^{-/-}$ SN *vs* TCL1 SN |
|---|---|---|---|---|---|---|
| CCL1 | 2.57 ± 0.3 | 5.86 ± 3.5 | 5.41 ± 2.7 | - | - | - |
| CCL11 | 3.11 ± 0.1 | 3.51 ± 0.7 | 3.21 ± 0.7 | - | - | - |
| CCL17 | 33.16 ± 0.5 | 37.74 +/- 4.5 | 62.53 +/- 13.1 | - | **p<0.05* | **p<0.05* |
| **CCL22** | **116.65 ± 50.98** | **324.23 ± 227.4** | **1605.74 ± 756.0** | ***p<0.05** | ****p<0.01** | ***p<0.05** |
| **CCL24** | **106.54 ± 23.5** | **279.96 ± 147.0** | **555.44 ± 211.3** | ****p<0.01** | *****p<0.001** | ****p<0.01** |
| CCL27 | 492.53 ± 90.3 | 445.98 ± 32.8 | 912.95 ± 365.5 | - | **p<0.05* | **p<0.05* |
| CXCL10 | 161.94 ± 6.6 | 315.49 ± 83.0 | 189.72 ± 48.68 | **p<0.05* | - | - |
| CXCL11 | 19.93 ± 0.3 | 17.50 ± 2.55 | 19.50 ± 1.3 | - | - | - |
| CXCL13 | 247.50 ± 34.9 | 220.33 ± 74.9 | 256.45 ± 80.2 | - | - | - |
| CXCL16 | 29.34 ± 7.5 | 40.55 ± 27.0 | 22.58 ± 15.8 | - | - | - |
| CXCL5 | 62.87 ± 14.6 | 66.74 ± 15.1 | 69.97 ± 19.6 | - | - | - |
| Fractalkine | 4.89 ± 0.9 | ND | 6.32 ± 2.3 | ND | - | ND |
| GM-CSF | 0.28 ± 0.0 | 0.21 ± 0.1 | 0.55 ± 0.4 | - | - | - |
| IFN-γ | 20.47 ± 1.6 | 19.30 ± 6.0 | 17.05 ± 1.0 | - | - | - |
| **IL-10** | **105.81 ± 12.4** | **279.37 ± 102.5** | **544.88 ± 154.5** | ***p<0.05** | *****p<0.001** | ***p<0.05** |
| IL-12 p40 | 3.05 ± 1.8 | 6.06 ± 5.0 | 1.68 ± 1.4 | - | - | - |
| IL-12 p70 | 0.13 ± 0.05 | 0.15 ± 0.06 | 0.13 ± 0.04 | - | - | - |

| | | | | | | |
|---|---|---|---|---|---|---|
| IL-13 | 0.35 ± 0.18 | 1.37 ± 0.85 | 12.03 ± 7.4 | - | **p<0.01 | *p<0.05 |
| IL-16 | 1113.91 ± 119.5 | 453.87 ± 257.6 | 561.02 ± 194.4 | *p<0.05 | - | - |
| IL-17A | 0.03 ± 0.001 | 0.03 ± 0.002 | 0.02 ± 0.001 | - | - | - |
| IL-1α | 0.0028 ± 0.0004 | 0.0075 ± 0.003 | 0.005 ± 0.0008 | **p<0.01 | - | - |
| IL-1β | 5.56 ± 0.9 | 2.0 ± 0.0 | 4.01 ± 2.8 | - | - | - |
| IL-2 | 3.83 ± 0.3 | 3.28 ± 0.6 | 4.61 ± 0.9 | - | - | - |
| IL-3 | ND | ND | ND | ND | ND | ND |
| IL-31 | 1.72 ± 0.87 | 1.22 ± 0.68 | 1.23 ± 0.67 | - | - | - |
| IL-4 | 10.26 ± 1.9 | 8.21 ± 1.1 | 11.68 ± 1.0 | - | - | - |
| IL-5 | ND | ND | ND | ND | ND | ND |
| IL-6 | 11.87 ± 1.0 | 7.16 ± 2.7 | 9.94 ± 5.0 | - | - | - |
| IL-9 | 161.01 ± 22.8 | 600.28 ± 176.3** | 890.48 ± 205.2 | **p<0.01 | ***p<0.001 | **p<0.01 |
| KC/CXCL1 | 8.48 ± 1.4 | 4.214 ± 1.1 | 5.23 ± 3.6 | *p<0.05 | - | - |
| MCP-1 | 1.1 ± 0.3 | 12.76 ± 11.1 | 5.2 ± 3.3 | - | - | - |
| MCP-3 | 5.68 ± 0.5 | 20.11 ± 5.0 | 29.52 ± 3.9 | *p<0.05 | *p<0.05 | - |
| MCP-5 | ND | 10.40 ± 2.9 | 4.03 ± 0.0 | ND | ND | **p<0.01 |
| MIP-1α | 17.43 ± 5.0 | 147.41 ± 25.6 | 221.76 ± 50.4 | **p<0.01 | ***p<0.001 | **p<0.01 |
| MIP-1β | 37.70 ± 11.3 | 116.66 ± 23.0 | 156.93 ± 55.0 | **p<0.01 | **p<0.01 | *p<0.05 |
| MIP-3α | 6.48 ± 0.5 | 5.26 ± 1.3 | 7.35 ± 2.8 | - | - | - |
| MIP-3β | 87.04 ± 13.1 | 98.754 ± 51.9 | 87.148 ± 27.7 | - | - | - |
| RANTES | 73.92 ± 23.3 | 106.97 ± 85.4 | 223.28 ± 124.15 | *p<0.05 | **p<0.01 | *p<0.05 |
| SDF-1α | 84.16 ± 34.3 | 283.48 ± 91.6* | 401.05 ± 111.7*/** | *p<0.05 | *p<0.05 | **p<0.01 |
| TNF-α | 16.4 ± 0.7 | 14.05 ± 5.2 | 15.06 ± 3.9 | - | - | - |

**Supplementary Table 2. List of soluble molecules whose expression is significantly modulated in leukemic cells from Eμ-TCL1/p66Shc$^{-/-}$ vs Eμ-TCL1 mice.** Fold change and *p* value of soluble factor expression extrapolated from Affymetrix array analysis showing differential expression patterns between leukemic Eμ-TCL1 (TCL1, n = 3) and Eμ-TCL1/p66Shc$^{-/-}$ (TCL1/p66$^{-/-}$, n = 3) cells. Differential expression criteria: *p*-value < 0.05, estimated fold change > 2. Differentially expressed genes are highlighted in bold.

| Protein name | Gene symbol | Ref Seq | *p*-value (TCL1-P66$^{-/-}$ *vs* TCL1) | fold-change (TCL1-P66$^{-/-}$ *vs* TCL1) |
|---|---|---|---|---|
| **Melanocyte protein** | *pmel* | NM_021882 | 0.00226604 | 164.968 |
| **Interleukin 9** | ***il9*** | **NM_008373** | **0.00263432** | **158.872** |
| **Osteocrin** | *ostn* | NM_198112 | 0.0078197 | 161.911 |
| **Defensin beta 8** | *defb8* | NM_153108 | 0.00971031 | -156.532 |
| **Apolipoprotein E** | *apoe* | NM_009696 | 0.0174238 | 22.271 |
| **C-type lectin domain family 2 member G** | *clec2g* | NM_001168223 | 0.0210042 | 160.048 |
| **Dystroglycan 1** | *dag1* | NM_001276481 | 0.0283007 | -192.572 |
| **Inactive serine protease 39** | *prss39* | NM_009355 | 0.0320975 | 152.112 |
| **85/88 kDa calcium-independent phospholipase A2** | *pla2g6* | NM_001199023 | 0.0330742 | 200.803 |
| **Pancreatic triacylglycerol lipase** | *pnlip* | NM_026925 | 0.0351441 | 175.732 |
| **Collagen alpha-1 (XVIII) chain** | *col18a1* | NM_001109991 | 0.035647 | -151.152 |
| **Immunoglobulin J chain** | *jchain* | NM_152839 | 0.0398578 | 174.997 |
| **C-C motif chemokine 22** | ***ccl22*** | **NM_009137** | **0.0467757** | **65.323** |

**Supplementary Table 3. mRNA levels of p66Shc and IL-9 in CLL cells inversely correlate with PD-1 surface expression in patient-matched $CD8^+$ cells.**

| Patient | p66Shc mRNA in CLL cells (ΔΔCt ratio) | IL-9 mRNA in CLL cells (ΔΔCt ratio) | Surface PD-1 in $CD8^+$ cells (MFI) | *IGHV* status |
|---|---|---|---|---|
| **# CLL 1** | 0.89 | 0.00 | 23.0 | M-CLL |
| **# CLL 2** | 0.64 | 0.00 | 21.7 | M-CLL |
| **# CLL 3** | 0.50 | 0.00 | 30.0 | M-CLL |
| **# CLL 4** | 0.56 | 0.00 | 36.0 | M-CLL |
| **# CLL 5** | 0.15 | 0.40 | 92.0 | U-CLL |
| **# CLL 6** | 0.46 | 0.00 | 37.0 | U-CLL |
| **# CLL 7** | 0.68 | 0.00 | 22.0 | M-CLL |
| **# CLL 8** | 0.01 | 1.70 | 188.0 | U-CLL |
| **# CLL 9** | 0.75 | 0.00 | 21.0 | M-CLL |
| **# CLL 10** | 0.50 | 0.00 | 17.5 | M-CLL |
| **# CLL 11** | 0.24 | 0.30 | 24.4 | M-CLL |
| **# CLL 12** | 0.01 | 0.55 | 30.0 | U-CLL |
| **# CLL 13** | 0.09 | 1.56 | 51.0 | U-CLL |
| **# CLL 14** | 0.20 | 0.93 | 33.0 | U-CLL |
| **# CLL 15** | 0.35 | 0.67 | 27.0 | M-CLL |
| **# CLL 16** | 0.02 | 1.41 | 34.0 | U-CLL |
| **# CLL 17** | 0.34 | 0.80 | 31.0 | M-CLL |
| **# CLL 18** | 0.08 | 1.00 | 44.0 | U-CLL |

**Supplementary Table 4. Clinical parameters of 4 CLL patients subjected to second line Ibrutinib treatment*.**

| | | CLL patients | | | |
|---|---|---|---|---|---|
| | | **# 1** | **# 2** | **# 3** | **# 4** |
| ***IGHV status*** | | Unmutated | Mutated | Mutated | Unmutated |
| ***TP53 status*** | | Wild-type | Wild-type | Wild-type | Wild-type |
| ***TP53 status*** | | Deleted and mutated | Mutated | Wild-type | Mutated |
| ***Follow-up [months]*** | | 32 | 21 | 33 | 19 |
| ***WBC (n/mL)*** | Before Ibrutinib treatment | 103000 | 74000 | 8560 | 138000 |
| | Follow-up Ibrutinib treatment | 40000 | 28000 | 5780 | 8410 |
| ***Ly (n/μL)*** | Before Ibrutinib treatment | 93000 | 68000 | 3900 | 130000 |
| | Follow-up Ibrutinib treatment | 36000 | 23000 | 2100 | 2200 |
| ***Hb (g/L)*** | Before Ibrutinib treatment | 127 | 150 | 134 | 95 |
| | Follow-up Ibrutinib treatment | 133 | 151 | 134 | 119 |
| ***PLT (n/μL)*** | Before Ibrutinib treatment | 530000 | 95000 | 67000 | 119000 |
| | Follow-up Ibrutinib treatment | 331000 | 80000 | 130000 | 166000 |
| ***RAI Stage*** | Before Ibrutinib treatment | 3 | 3 | 4 | 3 |
| | Follow-up Ibrutinib treatment | 0 | 1 | 0 | 1 |

*see Supplementary Methods for first line treatments.

**Supplementary Table 5. List of antibodies and reagents used in this study.**

| Reagent | Host | Clone | Source | Cat. n. | Concentration/ dilution |
|---|---|---|---|---|---|
| **CFSE** | - | - | Thermo Fisher Scientific | C34554 | 1.5 µM |
| **Propidium Iodide** | - | - | Sigma-Aldrich | 537059 | 0.5 µg/ml |
| **Monensin** | - | - | BioLegend | 420701 | 2 µM |
| **Fc-Block** | rat | 2.4G2 | BD Biosciences | 553141 | 1:50 |
| **PerCP-Cy5.5 IgM antibody** | rat | RMM-1 | BioLegend | 406512 | 1:30 |
| **PE anti-mouse CD5 antibody** | rat | 53 - 7.3 | BD Bioscience | 553022 | 1:40 |
| **FITC anti-mouse CD19 antibody** | rat | 1D3 | BD Bioscience | 553785 | 1:40 |
| **PE anti-mouse CD8a (Ly-2) antibody** | rat | 53 – 6.7 | BD Bioscience | 553033 | 1:50 |
| **PE anti-human CD8a antibody** | mouse | RPA-T8 | BioLegend | 301008 | 1:50 |
| **APC anti-human CD107a/LAMP1 antibody** | mouse | H4A3 | BioLegend | 328619 | 1:160 |
| **Anti-PD1 antibody** | rabbit | - | Novus | NBP1-77277 | 1:200 |
| **FITC anti-mouse CD45.1 antibody** | - | A20 | BioLegend | 110705 | 1:50 |
| **FITC anti-human CD69 antibody** | - | FN50 | BioLegend | 310904 | 1:100 |
| **Pe anti-human CD25 antibody** | - | BC96 | BioLeged | 302606 | 1:50 |

| | | | | | |
|---|---|---|---|---|---|
| **PerCP/Cyanine5.5 anti-human CD152 (CTLA-4) Antibody** | - | L3D10 | BioLegend | 349927 | 1:50 |
| **FITC anti-human CD223 (LAG-3) Antibody** | - | 11C3C65 | BioLegend | 369307 | 1:50 |
| **FITC-Annexin V** | - | - | BioLegend | 640906 | 1:50 |
| **Anti-CD3ζ antibody** | mouse | 6B10.2 | SantaCruz | Sc-1239 | 1:200 |
| **Anti- pTyr antibody** | mouse | 4G10 | Cell Signaling | 8954S | 1:100 |
| **Anti-PCNT antibody** | rabbit | - | Abcam | 4448 | 1:200 |
| **Alexa Fluor 555 F-actin** | - | - | Invitrogen | A34055 | 1:100 |
| **Alexa Fluor anti-mouse 488** | goat | - | Thermo Fisher Scientific | A11001 | 1:80 |
| **Alexa Fluor anti-rabbit 555** | goat | - | Thermo Fisher Scientific | A21428 | 1:80 |
| **Alexa Fluor anti-rabbit 647** | goat | - | Thermo Fisher Scientific | A21236 | 1:400 |
| **Recombinant Mouse IL-9** | - | - | R&D Systems | 409-ML-010 | 0.5 ng/ml |
| **Recombinant Human IL-9** | - | - | R&D Systems | 209-ILB-010 | 20 ng/ml |
| **Recombinant Human IL-10 Protein** | - | - | R&D Systems | 217-ILB | 0.5 ng/ml |
| **Recombinant human IL-2** | - | - | Miltenyi Biotech | 130-097-745 | 50 Units/mL |
| **IgG2B, isotype control antibody** | Rat | 141945 | R&D Systems | MAB0061 | 0.1 ng/ml |
| **Mouse IL-9 antibody** | Rat | 222622 | R&D Systems | MAB4091-100 | 0.1 ng/ml |
| **Human IL-9 antibody** | Mouse | 795908 | R&D Systems | MAB2092 | 1 ng/ml |

| | | | | | |
|---|---|---|---|---|---|
| **Human IL-10 antibody** | Mouse | 948505 | R&D Systems | MAB9184 | 1 ng/ml |
| **Neutralizing Human anti-PD-1 antibody** | Rabbit | | R&D Systems | AF1086 | 7.5 μg/ml |
| **B-1a Cell Isolation Kit, mouse** | - | - | Miltenyi Biotech | 130-097-413 | - |
| **Dynabeads™ Untouched™ Mouse CD8 Cells Kit** | - | - | Invitrogen | 11417D | - |
| **RosetteSep Human CD8+ T Cell Enrichment Cocktail** | - | - | StemCell Technologies | 15063 | - |
| **RosetteSep Human B Cell Enrichment Cocktail** | - | - | StemCell Technologies | 15064 | - |
| **Dynabeads Human T-activator CD3/CD28** | - | - | Gibco | 11132D | - |
| **Ibrutinib** | - | - | R&D Systems | | 10 μM |

**Supplementary Table 6. List of primers used in this study**

| **Quantitative RT-PCR** | **Forward 5'-3'** | **Reverse 5'-3'** |
|---|---|---|
| **Human IL-9** | ACCAGACCATGCTTCAGTGA | TCTTCAGAAATGTCAGCGCG |
| **Human IL-10** | TGCCTTCAGCAGAGTGAAGA | GGTCTTGGTTCTCAGCTTGG |
| **Human p66Shc** | TCCGGAATGAGTCTCTGTCA | GAAGGAGCACAGGGTAGTGG |
| **Human CCL22** | ACTGCACTCCTGGTTGTCCT | CGGCACAGATCTCCTTATCC |
| **Human CCL24** | GGAGTGGGTCCAGAGGTACAT | CAGGTGGTTTGGTTGCCAG |
| **Human HPRT1** | AGATGGTCAAGGTCGCAAG | GTATTCATTATAGTCAAGGGCATATC |
| **Mouse PD-1** | CATGCCCAGGTACCTCAGTT | GAACCCAACTCCAGGACAGA |
| **Mouse p66Shc** | TGAGTTGGGAGAGCAGAGGT | CTCATTCCGAAGTGGGTTGT |
| **Mouse IL-9** | CTTGCCTGTTTTCCATCGGG | CACGGCACCAGGAAAGAAAA |
| **Mouse GAPDH** | AACGACCCCTTCATTGAC | TCCACGACATACTCAGCAC |
